# Rapid modeling of experimental molecular kinetics with simple electronic circuits instead of with complex differential equations

**DOI:** 10.1101/2022.05.11.491353

**Authors:** Yijie Deng, Douglas Raymond Beahm, Xinping Ran, Tanner G Riley, Rahul Sarpeshkar

## Abstract

Kinetic modeling has relied on using a tedious number of mathematical equations to describe molecular kinetics in interacting reactions. The long list of differential equations with associated abstract variables and parameters inevitably hinders readers’ easy understanding of the models. However, the mathematical equations describing the kinetics of biochemical reactions can be exactly mapped to the dynamics of voltages and currents in simple electronic circuits wherein voltages represent molecular concentrations and currents represent molecular fluxes. For example, we theoretically derive and experimentally verify accurate circuit models for Michaelis-Menten kinetics. Then, we show that such circuit models can be scaled via simple wiring among circuit motifs to represent more and arbitrarily complex reactions. Hence, we can directly map reaction networks to equivalent circuit schematics in a rapid, quantitatively accurate, and intuitive fashion without needing mathematical equations. We verify experimentally that these circuit models are quantitatively accurate. Examples include i) different mechanisms of competitive, noncompetitive, uncompetitive, and mixed enzyme inhibition, important for understanding pharmacokinetics; ii) product-feedback inhibition, common in biochemistry; iii) reversible reactions, important in many metabolic pathways; and iv) translation and transcription dynamics in a cell-free system, which brings insight into the functioning of all gene-protein networks. We envision that circuit modeling and simulation could become a powerful scientific communication language and tool for quantitative studies of kinetics in biology.

## 1. Introduction

Kinetic modeling has been a powerful tool for studying biological systems from simple enzymatic reactions to metabolic pathways, drug kinetics in hosts, gene circuits in synthetic biology, and host-pathogen interactions (Alves et al., 2006; Resat et al., 2009; Bevc et al., 2011; Eshtewy and Scholz, 2020; Néant et al., 2021). Modeling molecular kinetics can provide quantitative insights and mechanistic understandings of biological systems. However, kinetic modeling of biological processes relies on a substantial number of mathematical equations to describe even simple biochemical reactions. The heavy dependence on long and tedious differential equations hinders many biologists from appreciating and taking advantage of kinetic modeling as a powerful tool for studying biological questions. In particular, for many biological researchers, the long list of parameters and abstract terms that are used during the process of mathematical derivation are tedious and difficult to follow. In addition, it can be challenging to resolve complex, nonlinear, coupled differential equations that require sophisticated algorithms/programs including numerical approaches (Bevc et al., 2011) for simulating time-course kinetics.

However, ordinary differential equations (ODEs), commonly used to model biochemical reactions and processes, can be represented by simple electronic circuits (Sarpeshkar, 2010; Teo and Sarpeshkar, 2020) in a mathematically exact fashion. We can thus take advantage of electronic design software to design circuits *in silico* that represent the kinetics of the target system and then run simulations in software without the need to manufacture the physical circuits (Teo et al., 2019a; Teo and Sarpeshkar, 2020). Therefore, not only can we visualize all the math equations in one circuit but also solve them by just running simulations on the circuit. Using virtual electronic circuits enables one to do rapid kinetic modeling of biochemical reactions without deriving tedious differential equations. In addition, circuit simulation in electronic design software is able to provide accurate time-course dynamics, not just equilibrium solutions. Circuit software has built-in algorithms to automatically solve underlying equations represented by the circuits, which has evolved over 75+ years of circuit design for multiple forms of design and analysis (Sarpeshkar, 2010).

The overall mechanism of the simulation is that, given some preset parameters of a circuit, the voltage and current at any node of the circuit at any time are readily available upon simulation; these voltages and currents exactly represent the corresponding molecular concentrations and molecular reaction flux rates, respectively. For example, we have used electronic circuits to model and simulate complex biological processes including genetic circuits in synthetic biology (Daniel et al., 2013; Teo et al., 2015; Zeng et al., 2018); kinetics of microbial growth and energetics (Deng et al., 2021); tissue homeostasis (Teo et al., 2019b); and virus-host interactions (Beahm et al., 2021). However, there are gaps in biologists’ understanding of electronic circuits and the underlying mathematics; and, in their understanding of the analogy of circuit variables to reaction kinetic parameters. These gaps have prevented many researchers from understanding circuit models and using circuits to do kinetic modeling in practice.

Therefore, an important goal of this work is to illustrate how the mathematics describing the kinetics of biochemical reactions can be exactly mapped to electronic circuits; and, to demonstrate how to use such circuits to do rapid kinetic modeling without deriving math equations. We start with the basics of a simple resistor-capacitor (RC) circuit and its use in representing the dynamics of a simple biochemical reaction. We then illustrate how to use circuits to model an enzyme-substrate reaction that is characterized by Michaelis-Menten kinetics, one of the most fundamental processes in biology. In addition to theoretical derivations, we also validate our circuits by fitting circuit models to experimental data. Finally, we demonstrate the wide applicability of circuit models in several fundamental biochemical reactions, including enzyme inhibition, product feedback inhibition, reversible reactions, and regulated transcription and translation in a cell-free systems. These circuits are fundamental to building circuit models for complicated biological networks/pathways without using cumbersome math equations; mature circuit-simulation software can then automatically provide accurate solutions including the time-course kinetics of molecules.

## 2. Results

### 2.1 Mapping a basic chemical reaction to a simple electronic circuit

To help biologists understand the basic mathematics of electronic circuits, we first derive, step-by-step, the basics of a simple RC circuit that is foundational for kinetic modeling in biological systems. The RC circuit consists of a resistor (R), a capacitor (C), and an input current (*I*_in_) that is generated by a voltage-controlled current generator (a ‘transconductor’) that converts the input voltage (V_in_) into the current (*I*_in_) (Figure 1). In electronics, a current is denoted by *I* (measured in Amps, A), a voltage is denoted by V (measured in Volts, V), a resistor has a resistance of R (measured in Ohms, Ω), and a capacitor has a capacitance of C (measured in Farad, F = A*s/V). As shown in Figure1, the input current (*I*_in_) goes through the capacitor and the resistor; the voltage (V) on the capacitor and the resistor keeps increasing until it reaches a steady state wherein the capacitor is fully charged; thereafter, all the input current (*I*_in_) goes through the resistor. To describe how the voltage (V) changes over time, we first calculate three currents in the circuit as below:

**Figure 1.**
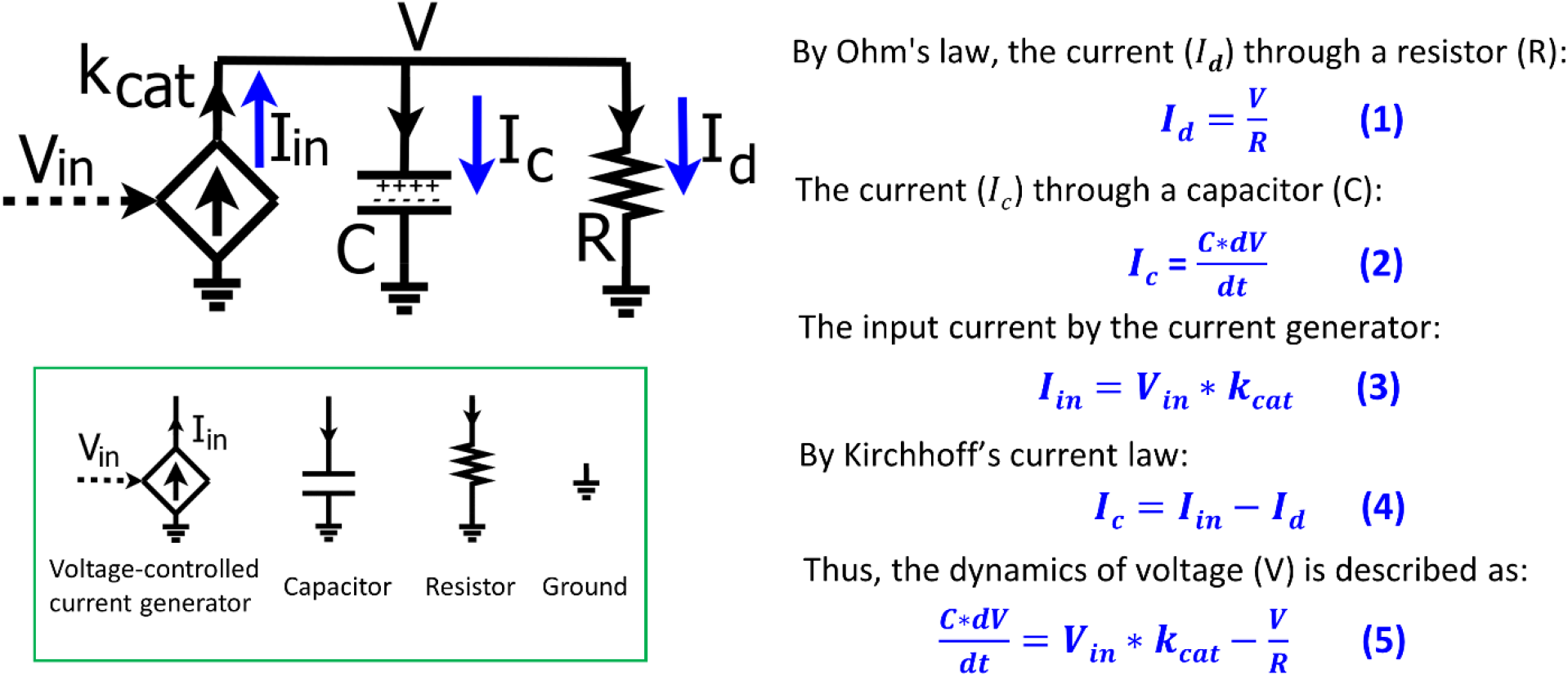
The basics of a resistor-capacitor (RC) circuit fed by a transconductor input. The input current is generated by the transconductor (diamond symbol), i.e., a voltage-controlled current generator that converts the input voltage (V_in_) into the input current (I_in_) with a conversion factor of k_cat_. The dynamics of the voltage (V) over the capacitor (C) and the resistor (R) are determined by the input current (I_in_) and the current (I_d_) through the resistor. Electronic circuit symbols are shown in the green box.

According to Ohm’s law, the current (*I*_*d*_) going through the resistor is defined by equation (1):

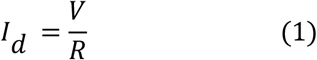

The total charge on the capacitor is Q and thus the current (*I*_*c*_) through the capacitor is described as:

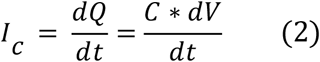

As mentioned above, the input current (*I*_*in*_) is generated by a voltage-controlled current source (the diamond-shaped device or ‘transconductor’ in Figure 1) that converts the input voltage (*V*_*in*_) into the current with a conversion factor (*k*_*cat*_), and thus we have:

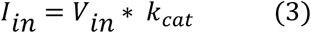

The input current is split into *I*_*c*_ and *I*_*d*_ in the circuit. By Kirchhoff’s current law, we have:

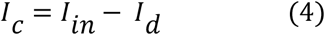

We substitute equations (1-3) into equation (4) and thus have equation (5) that describes the voltage dynamics in the RC circuit:

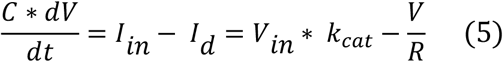

The dynamics of the voltage (V) over the capacitor and resistor are determined by the input current (*I*_*in*_) and the current through the resistor (*I*_*d*_). A simple analogy for the circuit is that the product concentration in a reaction system is determined by the production rate and the degradation rate. Given constant C and *k*_*cat*_ in equation (5), the voltage dynamics are thus determined by V_in_ and R which can be translated into biological relevance, as we discuss later. Equations (1-5) describe the basics of a simple RC circuit and are foundational for understanding the map between circuit modeling and mathematical modeling of kinetic processes. So, we have summarized their derivation in Figure 1. With the voltage dynamics described by equation (5), we normalize the equation by C such that the change of V over time is described as:

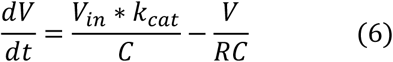

To simply the equation, we normally set C = 1 (F) = 1 (A*s/V) in circuit modeling and the above equation becomes:

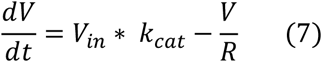

Equation (7) describes the voltage kinetics in the RC circuit (Figure 1) when C = 1. Setting C = 1 in circuit models enables a direct map to the mathematics behind the kinetics of a chemical reaction, as we illustrate in Figure 2. In Figure 2A, a substrate S is converted into a product P with a production rate constant of *k*_*cat*_ (1/s) and the product decays with a rate constant of 1/r (1/s). The kinetics of this reaction can be exactly described by an equivalent RC circuit fed by a transconductor (Figure 2B) which is identical to the RC circuit above (Figure 1). In the context of biological systems, we can consider the same reaction to be taking place in a container or a cell with a volume of C (liter, L) (Figure 2D). According to the law of mass action in chemistry, the production rate is proportional to the concentration of S and thus is *S* * *k*_*cat*_ (M/s); similarly, the decay rate of the product is P * (1/r) (M/s). The total amount of product P changes over time (mol/s) in the container, and is thus described as below:

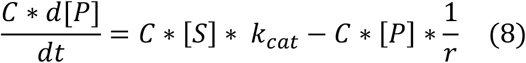

**Figure 2.**
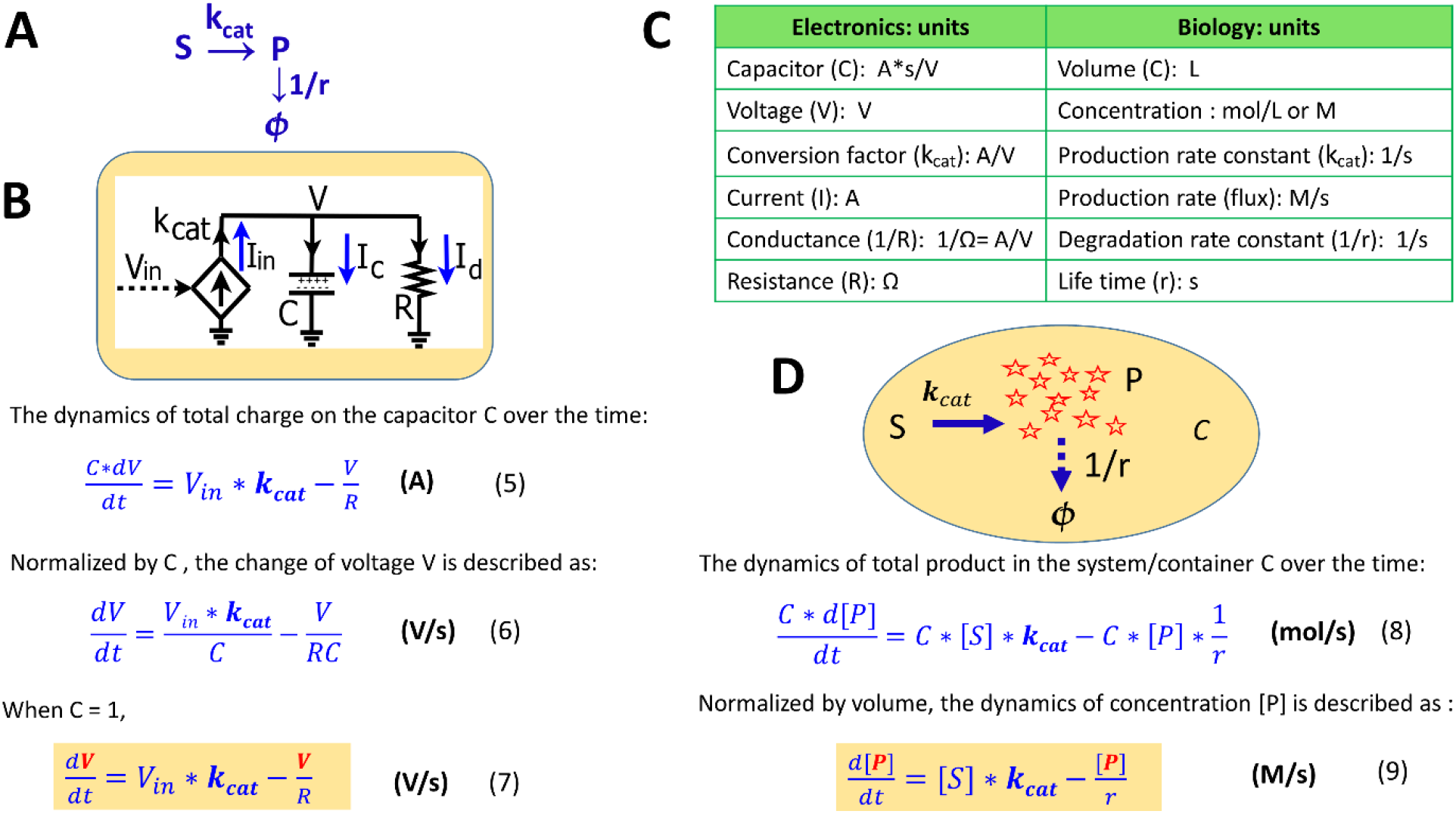
The mapping of an elementary biochemical reaction to an equivalent electronic circuit. (A) An example of a simple biochemical reaction wherein substrate S is converted to product P at a rate constant of *K*_*cat*_ (1/s) while the product also decays at a rate constant of 1/r (1/s). (B) A simple RC circuit in the context of the chemical reaction. (C) Translation of electronic variables into biochemical kinetics in a reaction. (D) The same biochemical reaction taking place in a container or a cell with a volume of C. The capacitance of a capacitor is normally set C = 1 A*s/V, which represents a volume-normalized container in a system (per L). Some important equations are summarized in this figure for comparison.

where [S] and [P] are the concentrations (M); C is the container volume (L); *k*_*cat*_ and 1/r are rate constants (1/s). Equation (5) is physically parallel to equation (8), wherein the former describes the change of the total charge of the capacitor while the latter describes the change of the total amount of product in a container. Then, we normalize equation (8) by the container volume C and thus have the concentration kinetics:

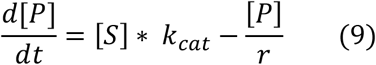

Equation (9) describes how the concentration of a product changes over time in a container, a cell, or in any reaction system. When comparing equation (9) to equation (7), we notice that they become mathematically identical with the input voltage V_in_ representing the substrate concentration [S]; the voltage V representing the production concentration [P]; *k*_*cat*_, the conversion factor for the current-generating transconductor, representing a production rate constant; and, the resistance R defining the time constant (or lifetime, r) of the product. The side-to-side comparison between electronic dynamics and chemical dynamics for this foundational production reaction is summarized in Figure 2C.

As in this production-reaction example, the dynamics in electronic circuits can be translated into the kinetics of biochemical reactions in more complex systems as well: the voltage (V) corresponds to the concentration (M) of a reagent in a chemical reaction; the current (A) is analogous to a reaction flux (M/s); the resistor R (Ω) defines degradation with 1/(RC) corresponding to a degradation rate constant (1/s) and RC corresponding to the equivalent time constant (lifetime); the capacitor with capacitance C = 1 A*s/V corresponds to a volume-normalized container in a biochemical system (per L). Unless otherwise mentioned, all capacitors in our circuits have C = 1. The dynamics of P are determined by the production flux and the degradation flux (Figure 2D) while the dynamics of the voltage (V) in the RC circuit (Figure 2B) are determined by the input source current (*I*_*in*_) and the sink current (*I*_*d*_).

### 2.2 Circuit modeling of Michaelis-Menten kinetics

We next demonstrate how to use circuits to simulate Michaelis-Menten kinetics for enzyme-substrate interactions. For a general enzymatic reaction (Figure 3A), enzyme E binds to a substrate S to form an intermediate enzyme-substrate complex ES at a rate constant of *k*_*f*_; ES will then either dissociate into E and S with a reverse rate constant of *k*_*r*_ or be converted to product P and free enzyme *E*_*free*_ with a rate constant of *k*_*cat*_. From mass conservation, we have:

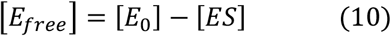

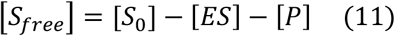

**Figure 3.**
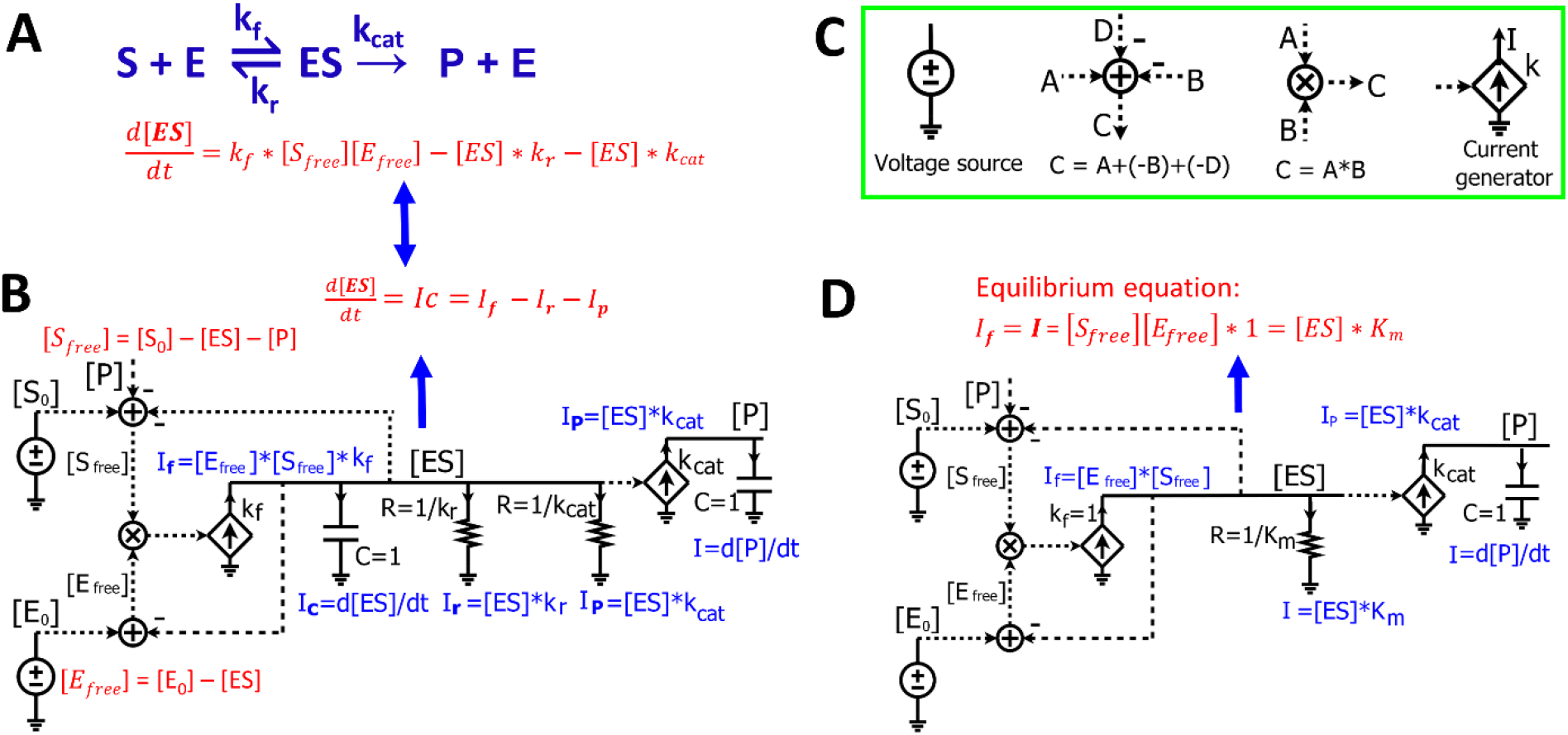
Modeling Michaelis-Menten kinetics of enzymatic reactions by simple electronic circuits. (A) A general enzymatic reaction wherein the enzyme E binds to the substrate S, forming an enzyme-substrate complex ES, which converts S to a product P. (B) The electronic circuit exactly describes the kinetics of the enzymatic reaction in (A). All the math equations describing the voltages and/or currents of the circuit are indicated near the corresponding nodes. The dashed lines are wires connecting the same voltage between two nodes/components in the circuit and have no current running through them. The voltages labeled with the same names indicate that they have the same values. The voltages are mainly for math calculations such as calculating the mass conservation of a reagent via the adder/subtracter blocks, or multiplying of two concentrations via a multiplier block. They are also used as inputs to voltage-dependent current generators (transconductors, the diamond symbols) to control their output currents. (C) Electronic symbols used in the circuits in addition to the symbols from Figure 1. (D) The Michaelis-Menten circuit of (A), but with a steady-state approximation such that the [ES] capacitor is removed. Since the capacitor has been removed, resistors are directly related to steady-state Michaelis-Menten constants only and do not affect dynamic parameters like time constants. The resistor R = 1/K_m_ (Ω) and K_m_ is in the standard molar concentration unit, M.

where [E_0_] and [S_0_] are the initial concentrations of enzyme and substrate, respectively. The enzyme-substrate complex is converted into product P with a rate proportional to its concentration [ES], such that:

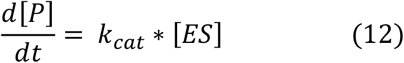

The kinetics of [ES] is determined by three fluxes: the forward reaction rate, *k*_*f*_ * [*E*_*free*_][*S*_*free*_], the reversed reaction rate, [*ES*] * *k*_*r*_, and the catalytic reaction rate [*ES*] * *kcat*. Therefore, the dynamics of [ES] are described as below:

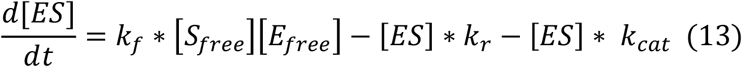

The circuit (Figure 3B) exactly represents the enzymatic reaction (Figure 3A). In this circuit, voltages of the wires are labeled with names corresponding to components of the enzymatic reaction. The dashed lines are wires that don’t have current going through them but still have the same voltage as the wires or nodes that they originate from. We first derive equations for currents and voltages in the circuit. Since there is no current running through any of the dashed lines/wires, this circuit (Figure 3B) is similar to the circuits we derived above (Figures 1 and 2B), consisting of two RC blocks connected together, one with two resistors and the other with no resistors. The dynamics of the voltage [ES] are determined by three currents: *I*_*f*_, *I*_*r*_ and *I*_*p*_. Because the voltage across the resistors and capacitor is [ES], by Ohm’s law, the current through the resistor (R =1/*k*_*r*_) is: *I*_*r*_ = [ES]/R = [ES]* *k*_*r*_ which represents the reverse reaction rate/flux; similarly, given the other resistor (R = 1/*k*_*cat*_), the current through it is: *I*_*p*_ = [*ES*] * *k*_*cat*_ which represents the catalytic flux; given that the current generator has a conversion factor of *k*_*f*_ and an input voltage [*E*_*free*_] * [*S*_*free*_] that is calculated by the multiplier, we have the input current *I*_*f*_ = [*E*_*free*_] * [*S*_*free*_] * *k*_*f*_. We note that the latter three currents are mathematically exactly the same as the three reaction fluxes in the enzymatic reaction (equation 13). Finally, the current through the capacitor is 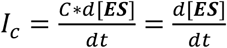 (when C=1) (Figure 3B). According to Kirchhoff’s current law, the current through the capacitor is given by:

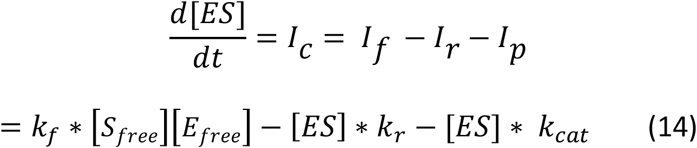

Equation (14) describes the dynamics of the voltage [ES] in the RC circuit (Figure 3B), which is exactly the same as equation (13) that we derived from the enzymatic reaction (Figure 3A). Just as in the enzymatic reaction, the dynamics of [ES] are determined by one generation reaction flux and two consumption fluxes (equation 13); the dynamics of the voltage [ES] in the RC circuit are determined by one source generation current and two sink consumption currents through the resistors (equation 14).

Finally, the voltage [P] is determined by *Ip* which is generated by a current generator with an input voltage [ES] and a conversion factor of *k*_*cat*_. Since all current generated flows into the capacitor (C = 1), we have:

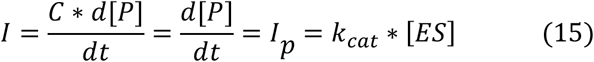

Equation (15) describes the dynamics of the voltage [P] in the RC circuit, which is exactly the same as equation (12) that describes the product kinetics in the enzymatic reaction.

In the circuit model (Figure 3B), two adders are used to calculate [*E*_*free*_] and [*S*_*free*_] based on the law of mass conservation. The binding of enzyme and substrate is represented by the multiplier symbol resulting in a signal, [*E*_*free*_] * [*S*_*free*_], which is the input voltage used to generate current in the first transconductor in Figure 3B. Given the conversion factor of *k*_*f*_, the resulting current is [*E*_*free*_] * [*S*_*free*_] * *k*_*f*_.

Since all the equations describing the electronic circuit and the enzymatic reaction (Figures 3 A,B) are mathematically identical, we can directly use the electronic circuit to simulate the kinetics of enzymatic reactions without deriving the underlying equations. The changes of concentrations and reaction fluxes over time are directly mapped to the corresponding changes in voltages and currents, respectively. Therefore, electronic circuits enable a powerful and intuitive method for visualizing multiple math equations in one pictorial schematic. Using these circuits is especially advantageous when one wants to simulate complicated biological pathways/networks where hundreds of differential equations can be represented in a single circuit. To draw/construct and simulate such electronic circuits, multiple electronic software packages are widely and easily available, including Cadence (Cadence Design Systems, Inc.), CircuitLab (https://www.circuitlab.com/), or MATLAB Simulink/Simscape Electrical (The MathWorks, Inc.). Once the circuits are constructed, we can simply run simulations with these tools. The dynamics of the voltages and currents then directly represent real-time changes of the concentrations and reaction fluxes in biochemical reactions, respectively.

The circuit in Figure 3B is an exact circuit for representing biochemical reactions without using any mathematical assumptions/approximations; however, circuit simulation requires known values of *k*_*f*_ and *k*_*r*_ which are not normally available for most enzymatic reactions. To circumvent this requirement and make the circuit more useful in practice, we apply the same assumptions that Michaelis-Menten equation uses to simplify dynamics: Under the quasi steady-state assumption that enzyme-substrate binding is much faster than the substrate-to-product conversion output reaction, the ES concentration is assumed to reach a steady state almost instantaneously. Therefore,

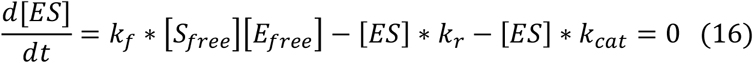

Normalizing the equation by *k*_*f*_ and grouping the [ES] terms, we have:

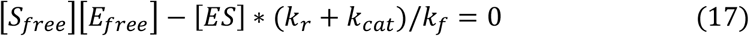

Letting *K*_*m*_ = (*k*_*r*_ + *k*_*cat*_)/*k*_*f*_, we have the equilibrium equation below:

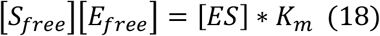

where *K*_*m*_ (M) is the Michaelis-Menten constant. Accordingly, we can also modify the circuit of Figure 3B to reflect the equilibrium equation (18). Since it is assumed that [ES] reaches a steady state instantaneously, it means that the capacitor connected to the [ES] voltage node is zero (Figure 3B), architecting an effective RC time constant of zero. Accordingly, we remove the capacitor for [ES] from the circuit in Figure 3B. Next, to normalize both forward and reverse reaction rates by *k*_*f*_, we set *k*_*f*_ = 1 in the circuit. Finally, since we have combined *k*_*f*_ and *k*_*cat*_ as in the equation (17), we merge the two currents, *I*_*r*_ and *I*_*p*_, into one current ([*ES*] * (*k*_*r*_ + *k*_*cat*_)/*k*_*f*_) through a resistor with resistance of 1/*K*_*m*_. Therefore, we have a simplified circuit (Figure 3D) containing only the parameters *K*_*m*_, *k*_*cat*_, S_0_, and E_0_, which are readily available from experiments.

We can easily confirm that the simplified circuit correctly reflects equation (18); the forward current generated by the current generator is *I*_*f*_ = [*S*_*free*_][*E*_*free*_], and the current that goes into the resistor is: *I* = [*ES*] * *K*_*m*_. These two currents are equal since all generated current goes through the resistor. Therefore, we have *I*_*f*_ = *I* = [*S*_*free*_][*E*_*free*_] = [*ES*] * *K*_*m*_, which is equation (18) describing [ES] in the reaction. This simplified circuit (Figure 3D) exactly characterizes the kinetics of the enzymatic reaction (Figure 3A) under the steady-state approximation and is the Michaelis-Menten equation in the circuit form. This Michaelis-Menten circuit is the basic building block for enzymatic reactions and can be easily extended into more complicated circuits for different mechanisms of enzyme inhibition as we show below. It should be noted that besides ensuring enzyme conservation, i.e., that [*E*_*free*_] and [*ES*] sum to [*E*_0_], the circuit of Figure 3D also ensures substrate conservation: [*S*_*free*_] = [*S*_0_] − [*ES*]. Ensuring conservation of both the enzyme and substrate species allows the Michaelis-Menten circuit to be more robust and accurate, especially under scenarios where the enzyme concentration and substrate concentration are comparable (discussed later).

### 2.3 Circuit modeling of a hydrolytic reaction by beta-galactosidase

To validate the Michaelis-Menten circuit (Figure 3D), we fit our circuit model to experimental data that we collected from an enzyme-substrate reaction. We chose to use a hydrolytic reaction wherein beta-galactosidase is the enzyme and ONPG is the substrate (Figure 4A). We ran the circuit simulation with the four necessary parameters (*K*_*m*_, *k*_*cat*_, S_0_ and E_0_), which were all experimentally determined under our test conditions. As we mentioned above, the reaction flux is given by the current *I*_*p*_ in the circuit while the voltage [S_0_] reflects the initial substrate concentration. As expected, the circuit model perfectly matches the measured initial reaction rates when we varied the initial substrate concentrations (Figure 4B). The predicted curve (V_0_ ∼ S_0_) is a classic hyperbolic curve for an enzymatic reaction; as S_0_ increases the initial reaction rate also increases until it reaches the maximal rate. For Lineweaver-Burk plotting, the circuit model accurately predicts a straight line (1/V_0_ versus 1/S_0_) with K_m_ = 0.167 mM (derived from the X-intercept in Figure 4C) and V_max_ = 0.00087 mM/s (derived from the Y-intercept in Figure 4C) with an excellent fit to the experimental data.

**Figure 4.**
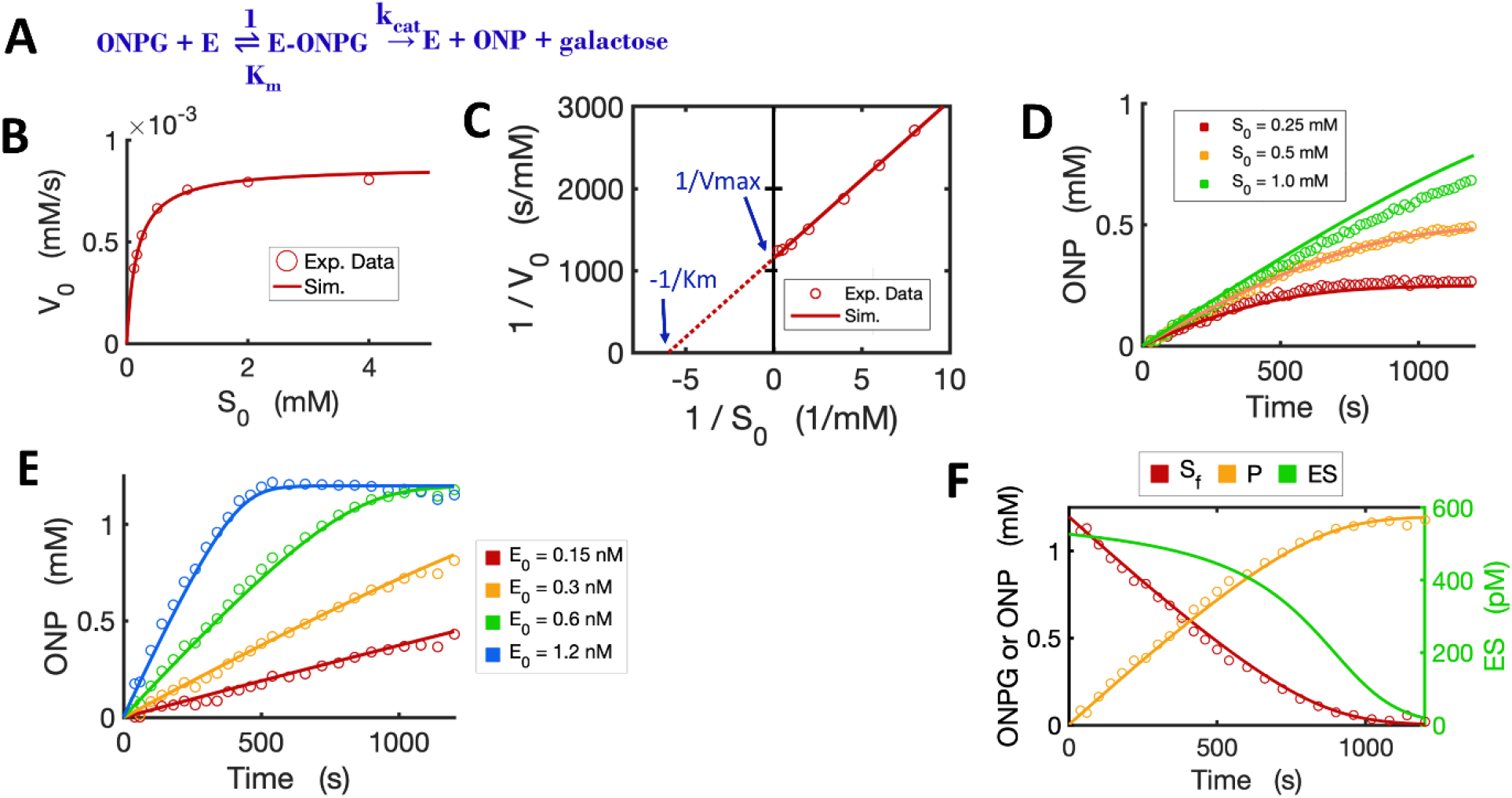
Circuit modeling of the kinetics of the beta-galactosidase reaction. (A) The enzymatic reaction scheme for E (beta-galactosidase) with its substrate ONPG. (B) Circuit simulation curve of initial substrate concentration [S_0_] versus initial reaction rate (V_0_) fitted to the experimental data (when E_0_ = 0.3 nM). (C) Lineweaver-Burk plot showing linearized curves of 1/S_0_ ∼1/V_0_ with a Y-intercept of 1/V_max_ and X-intercept of -1/K_m_. (D) Circuit simulation curves of product dynamics over time with varying initial S concentrations fitted to experimental data with E_0_ = 0.3 nM. (E) Circuit simulation curves of product dynamics over time with varying initial enzyme concentrations (0.15-1.2 nM) fitted to experimental data with S = 1.2 mM. (F) Model-predicted curves of the dynamics of [S_f_], [ES] and [P] over time with E_0_ = 0.6 nM and S_0_ = 1.2 mM. The data points for [P] are experimental data while the data points for the free substrate [S_f_] are calculated from stoichiometric conservation to be [S_f_] = [S_0_] -[P]. All simulation curves are obtained from the Michaelis-Menten circuit model with experimentally measured K_m_ = 0.167 mM and *K*_*cat*_ = 2903/s. Note that the resistor R = 1/K_m_ = 1/0.000167 = 5988 Ω. All data points are means of three independent replicates. The standard deviations are relatively small (less than 20% of the corresponding mean) and are not shown.

The circuit model can also accurately simulate the time-course dynamics. As shown in Figure 4D, the simulation curves of [P] dynamics fit our experimental data closely under varying initial substrate concentrations. In addition, when we changed the initial concentration of the enzyme with a constant substrate concentration, as expected, the circuit model predicts product dynamics that are in good agreement with our experimental data (Figure 4E). As more enzyme is added, the reaction consumes substrate faster and reaches a plateau earlier. The circuit model also accurately predicts the dynamics of [S_free_], [P], and [ES] over time (Figure 4F).

### 2.4 Circuit modeling of competitive inhibition and product-feedback inhibition

With the basic Michaelis-Menten circuit validated, we developed a circuit model for competitive inhibition. In the classic competitive inhibition model, an inhibitor binds to an enzyme at the substrate-binding site and competes with the substrate for the free enzyme, as shown in the reaction scheme (Figure 5A). To model the competitive inhibition in a circuit, we only need to add an enzyme-inhibitor binding circuit to the same Michaelis-Menten circuit (Figure 3D). Given the assumption that inhibitor binding is also much faster than the catalytic reaction and that the enzyme-inhibitor complex [*EI*] also, therefore, reaches steady-state instantaneously, we have the equilibrium condition:

**Figure 5.**
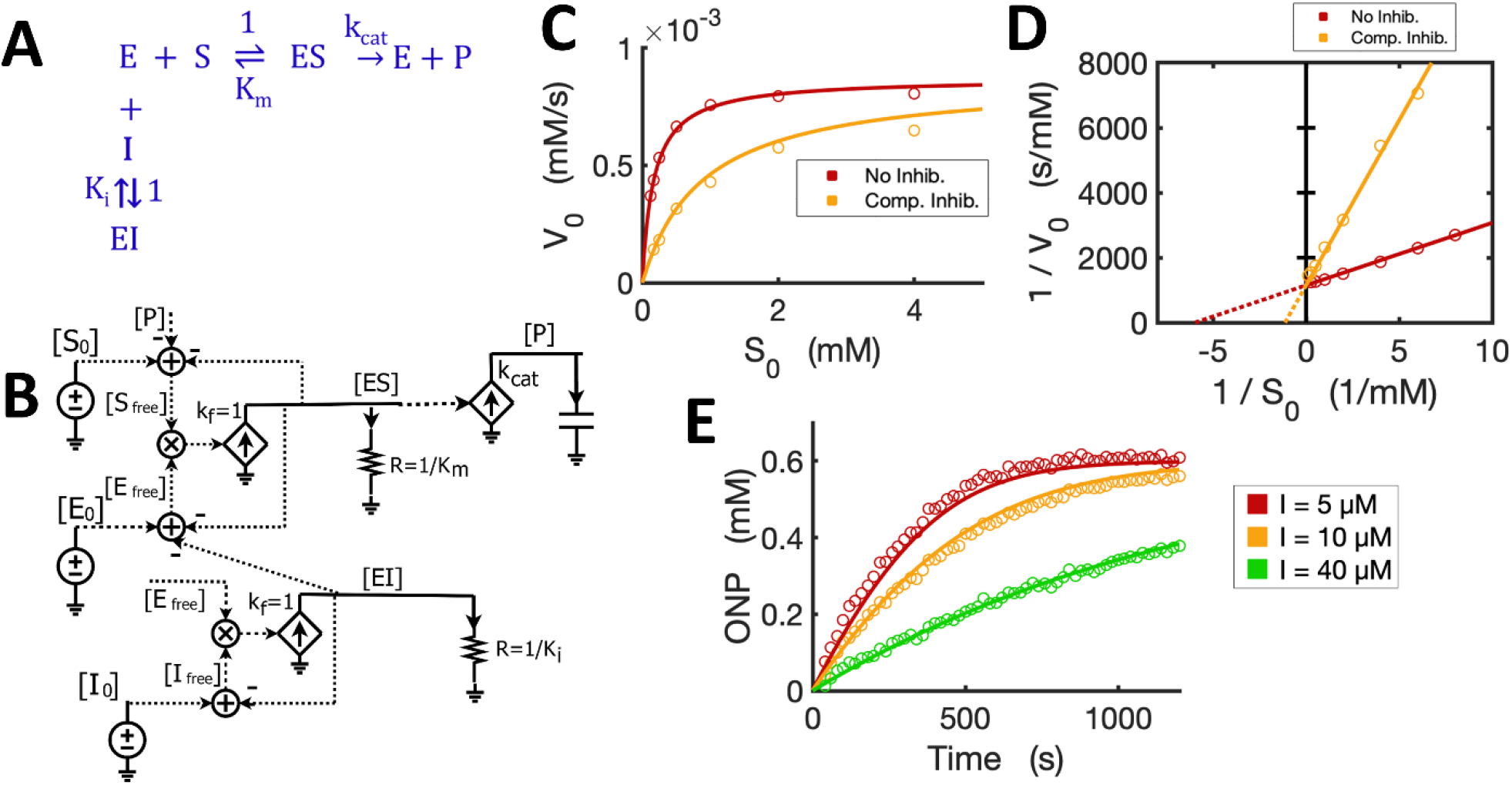
Circuit model of competitive inhibition. (A) The classic reaction scheme for competitive inhibition. (B) The circuit model for competitive inhibition. In this case, E is beta-galactosidase, S is ONPG, and the competitive inhibitor is phenylethyl beta-D-thiogalactopyranoside (PETG). (C) Simulation curves of initial substrate concentration S_0_ versus initial reaction rate V_0_ fitted to experimental data in the absence and presence of PETG (10 μM) when E_0_ = 0.3 nM. (D) Lineweaver-Burk plot showing linearized curves of 1/V_0_∼1/S_0_ fitted to experimental data with or without the inhibitor, which have the same Y-intercept, 1/V_max_. (E) The model curves of product dynamics over time with varying inhibitor concentrations fitted to experimental data points (E_0_ = 1.0 nM, S_0_ = 0.6 mM). All simulation curves are obtained from the circuit model with experimentally measured K_i_ = 2.33 μM, and the same K_m_ and *K*_*cat*_ from Figure 4. All data points are means of three independent replicates. The standard deviations are relatively small (< 20% of the corresponding mean) and are not shown.

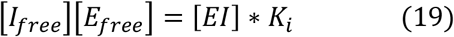

where *K*_*i*_ is the dissociation constant for the inhibitor. Equation (19) is similar to equation (18), so we make a similar circuit (the lower circuit of Figure 5B) representing the equilibrium enzyme-inhibitor binding as described in equation (19). In this circuit block, all we need is to set R = 1/K_i_ to reflect the inhibitor binding constant. The voltage [*EI*] in the circuit is then wired to the [E_0_] adder to account for the consumption of free enzyme that has been competitively bound by the inhibitor. Therefore, we have a competitive inhibition circuit model (Figure 5B). We used a competitive inhibitor of beta-galactosidase and experimentally demonstrated that this circuit is identical to the classic equations describing the kinetics of competitive inhibition. The circuit model accurately predicts the relationship between substrate [S_0_] and initial reaction rates (S_0_ ∼ V_0_) in the presence and absence of the inhibitor (Figure 5C). As [S_0_] increases, the initial reaction rate V_0_ also increases and eventually will reach the same maximal rate V_max_ even in the presence of the inhibitor. The linearized curves (Lineweaver-Burk plot) show the expected behavior of competitive inhibition where the inhibitor increases the apparent K_m_ but not the maximal reaction rate (V_max_) (Figure 5D). In addition, the circuit model can also exactly predict the product dynamics over time under different inhibitor concentrations (Figure 5E). Thus, using our experimental data, we have verified that the circuit model accurately describes the kinetics for competitive inhibition.

The circuit for competitive inhibition is a useful building block and can be used to construct circuits for complicated biological pathways when there are competitive inhibitors involved. As an example, we now use the competitive inhibition circuit (Figure 5B) to model the kinetics of product inhibition in an enzymatic reaction. Product inhibition is a common way to regulate reaction rates in metabolic pathways. Based on previous reports that beta-galactosidase can be competitively inhibited by a relatively high concentration of its own product galactose (Portaccio et al., 1998; Nguyen et al., 2006), we easily construct a reaction scheme for the product inhibition wherein the product galactose competes for free enzyme (Figure 6A): In Figure 5B, we simply replace the inhibitor I_0_ with product [Gal_0_] that can be externally added to the reaction and also wire newly produced [Gal] to the total product pool to architect the feedback inhibition; the resultant circuit is shown in Figure 6B.

**Figure 6.**
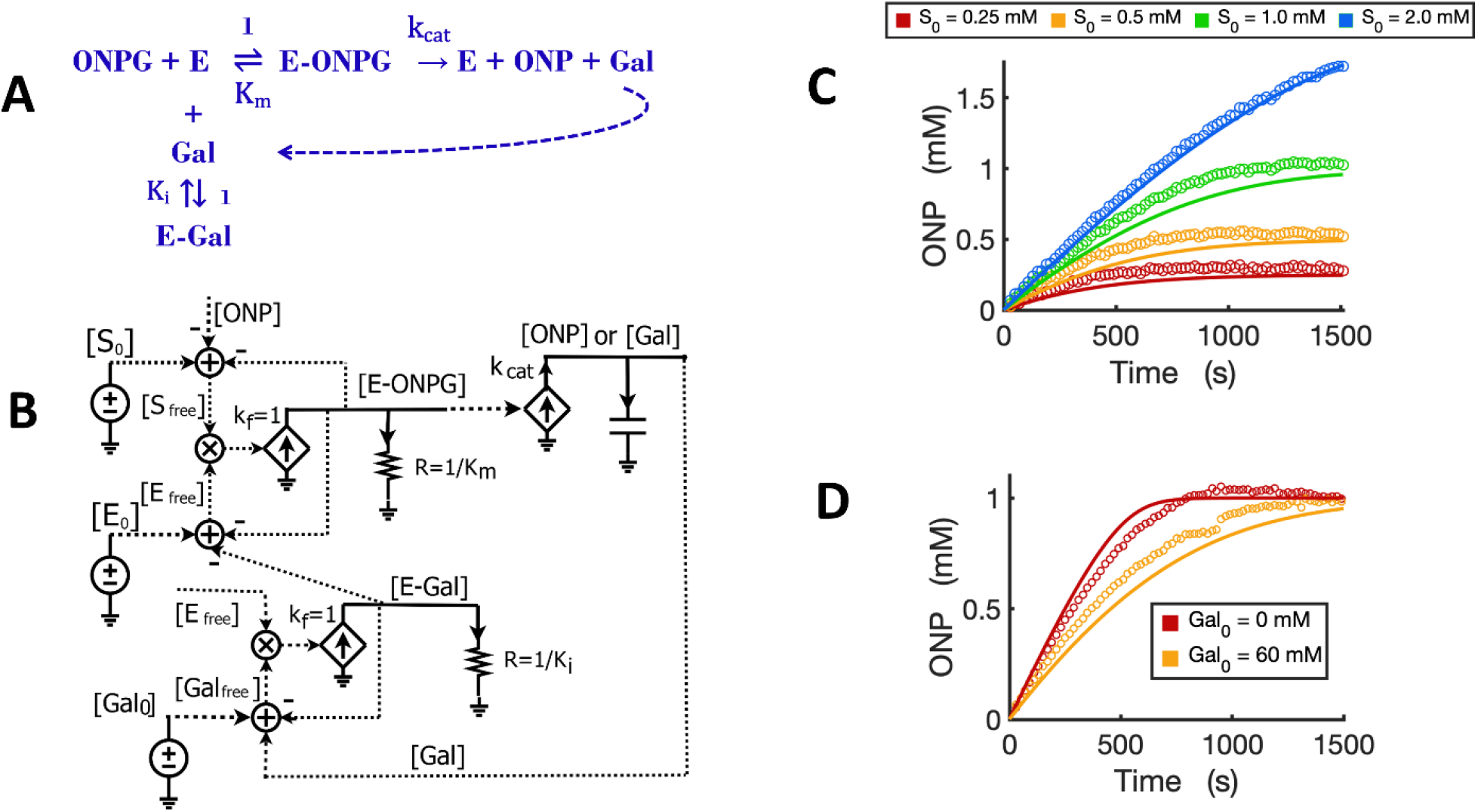
Circuit modeling of product feedback inhibition. (A) The scheme of product feedback inhibition based on the competitive inhibition. Galactose [Gal] is one of the products and is also an inhibitor to the enzyme, beta-galactosidase. (B) The circuit model for product feedback inhibition: [S_0_] is the initial ONPG added to the reaction while [Gal_0_] is the initial galactose added to the reaction. (C) Model curves of product dynamics over time fitted to experimental data with varying initial substrate concentrations (when [Gal_0_] = 40 mM, E_0_ = 0.7 nM). (D) Model curves of product dynamics fitted to experimental data with or without galactose added to the reaction (when ONPG = 1mM, E_0_ = 0.85 nM). All simulation curves are obtained from the circuit model (B) with experimentally measured K_i_ = 13.7 mM, and the same K_m_ and *K*_*cat*_ from Figure 4. All data points are means of three independent replicates with standard deviations less than 20% of the corresponding mean (not shown).

It is worth noting that simple rewiring and reuse of circuit building blocks avoids the need for any math equations, and preserves physical and chemical intuition. We can directly and rapidly map the reaction mechanism of Figure 6A to a quantitatively accurate representation of its function and dynamics in Figure 6B. The implicit (caused by subtractive inputs from the “use-it-and-lose-it” mass conservation in Figures 4-6) and explicit (due to product inhibition) feedback loops are all evident and clearly represented.

We validated the product-inhibition circuit model by fitting it to experimental data. The circuit model shows good fits to the experimental data for product dynamics over time when varying substrate concentrations were added but with constant galactose concentration (Figure 6C). In another experiment, we compared the reaction with or without the product galactose added before starting the reaction. As expected, when the initial amount of galactose is added, the reaction is inhibited and takes a longer time to reach a plateau wherein all substrate has been consumed (Figure 6D).

### 2.5 A generalized circuit block for enzyme inhibition

We next sought to develop a generalized circuit model for all types of enzyme inhibition including competitive, non-competitive, and uncompetitive inhibition. In the generalized reaction scheme (Figure 7A), the enzyme forms ES, EI, and ESI complexes with the substrate, inhibitor, or both, respectively; the specific reaction fluxes can be derived from the corresponding rate constants. We can directly translate the reaction scheme into an equivalent circuit (Figure 7B) that exactly describes all the dynamics of each species in the reaction and has identical math equations, as we derive below. Based on the mass conservation law, we have the following relationships for [*S*_0_], [*E*_0_] and [*I*_0_] (Figure 7A):

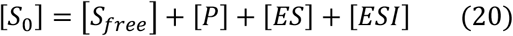

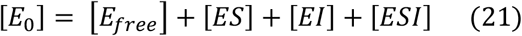

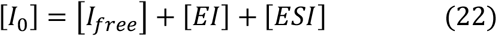

**Figure 7.**
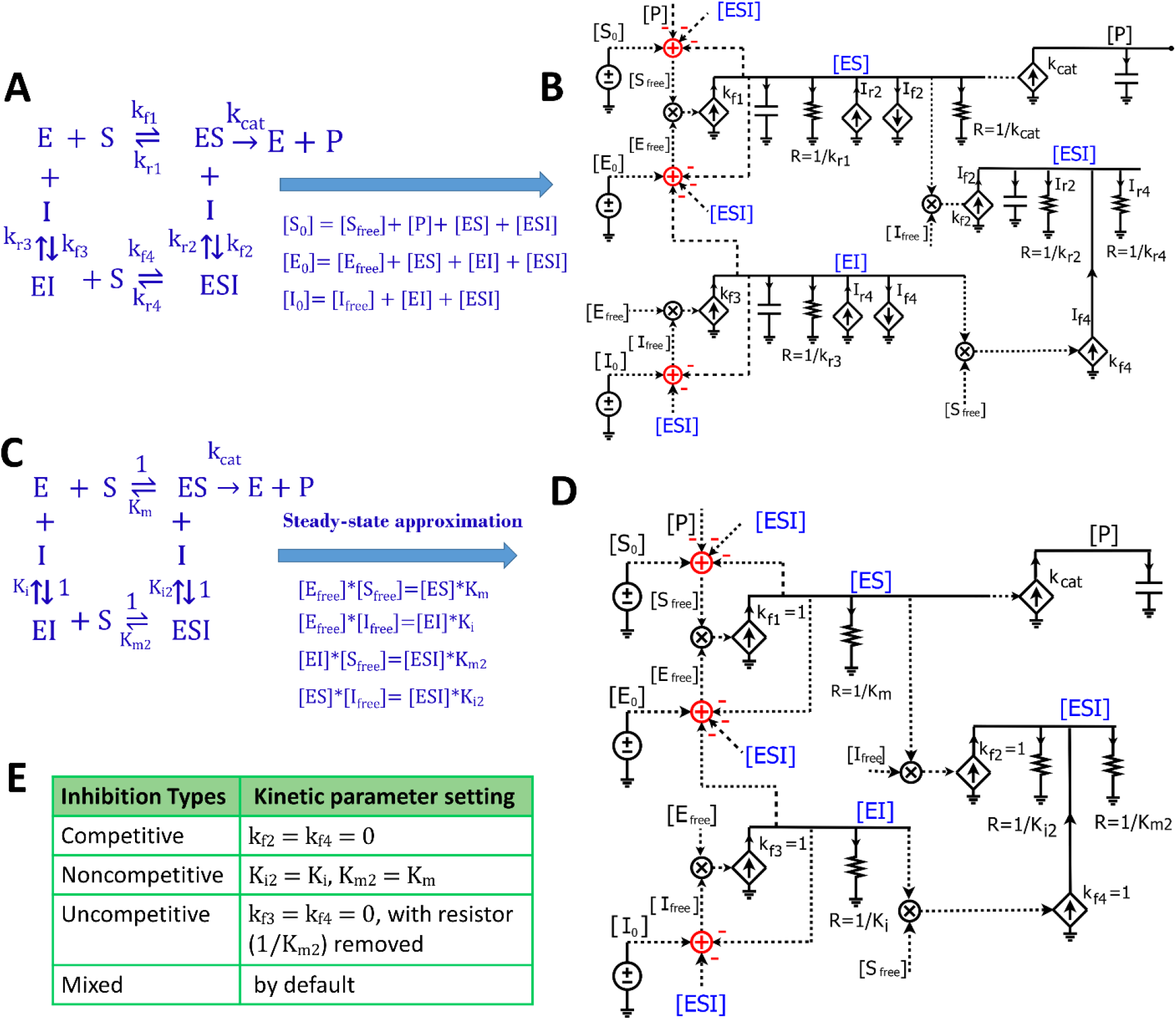
Generalized circuit models for enzyme inhibition. (A) A reaction scheme of general inhibition using rate constants without any approximation or assumption. (B) A circuit model translated from the reaction scheme in (A) describes reaction dynamics. All symbols used in this circuit are the same as the ones used in Figures 1 and 3. The dependent current generators (diamond symbols) can provide input source currents (arrow up) or sink currents (arrow down) at nodes that they are wired to. Voltages or currents labeled with same names indicate that they have the same values: in accord with the reaction, the same current or voltage is appropriately re-used or regenerated at multiple locations with the use of implicit rather than explicit wiring to avoid clutter. (C) The reaction scheme of general inhibition in (A), but with a steady-state approximation (all complexes reach equilibria instantaneously). Here, normalized k_f_ parameters are set to 1, such that all k_r_ parameters are mapped to their corresponding equilibria dissociation-constant (K_m_ or K_i_) values. (D) The generalized circuit for enzyme inhibition translated from the reaction scheme in (C). Note that, in accord with the steady-state approximation, the capacitors in (B) are removed in (D); and kinetic parameters in the reaction scheme (C) are mapped to equivalent circuit parameters in (D). The latter circuit can simulate all common enzyme-inhibition mechanisms including competitive, noncompetitive, uncompetitive, and mixed inhibition. (E) Kinetic parameter settings for different types of enzyme inhibition.

Such mass conservation is represented in the equivalent circuit (Figure 7B) by an adder block with a positive conserved total species input (S_0_, E_0_, or I_0_); subtractive (negative) inputs are caused by the use of the species (for binding or product transformation) to generate other species (P, ES, EI, or ESI); finally, free variables (S_free_, E_free_, or I_free_) are the resulting outputs of the adder blocks. The subtractive inputs always cause the ‘use-it-and-lose-it’ implicit negative-feedback loops in chemical reaction networks (Teo et al., 2015; Teo and Sarpeshkar, 2020). As shown in Figure 7A, the dynamics of [*ES*] are determined by five reaction fluxes, including two generation fluxes and three consumption fluxes. Therefore, we have:

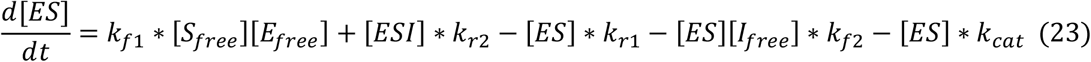

Likewise, the voltage dynamics of [*ES*] in the circuit (Figure 7B) are also determined by five currents/fluxes. The two generation fluxes include the forward reaction flux, *k*_*f*1_ * [*S*_*free*_][*E*_*free*_] for enzyme-substrate binding indicated by the *k*_*f*1_ current source, and the reverse reaction flux (*I*_*r*2_), [*ESI*] * *k*_*r*2_ from [*ESI*], indicated by the *I*_*r*2_ current source; the three consumption fluxes include the dissociation reaction flux, [*ES*] * *k*_*r*1_, indicated by the 1/*k*_*r*1_ resistor, the reaction flux for ES and inhibitor binding (*I*_*f*2_), [*ES*][*I*_*free*_] * *k*_*f*2_, indicated by the *I*_*f*2_ sink current source, and the catalytic reaction flux, [*ES*] * *k*_*cat*_.

The dynamics of [*EI*] are determined by two generation fluxes and two consumption fluxes in the reaction scheme (Figure 7A). Therefore, we have:

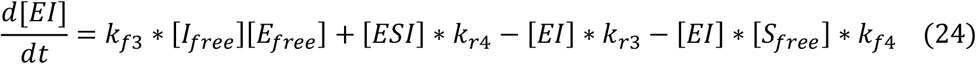

Similarly, in the circuit (Figure 7B), the voltage dynamics of [*EI*] are also determined by two source currents and two sink currents. Two supply currents include *k*_*f*3_ * [*I*_*free*_][*E*_*free*_], indicated by the *k*_*f*3_ current source, and the dissociation flux (*I*_*r*4_), [*ESI*] * *k*_*r*4_, indicated by the *I*_*r*4_ current source. The two sink currents include the current through the resistor (1/*k*_*r*3_), [*EI*] * *k*_*r*3_, and the current through the *I*_*f*4_ current source, which is [*EI*] * [*S*_*free*_] * *k*_*f*4_.

The dynamics of [*ESI*] in the reaction scheme (Figure 7A) are quantified by two generation fluxes and two consumption fluxes, so we have:

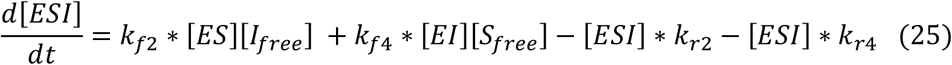

This equation is also exactly reflected by the voltage dynamics of [*ESI*] in the circuit (Figure 7B), which are determined by two source currents and two sink currents. As shown in the circuit, the two source currents include one from the *k*_*f*2_ current source, *k*_*f*2_ * [*ES*][*I*_*free*_], and one from the *k*_*f*4_ current source, *k*_*f*4_ * [*EI*][*S*_*free*_]. The two sink currents are indicated by two resistors 1/*k*_*r*2_ and 1/*k*_*r*4_, with quantities of [*ESI*] * *k*_*r*2_, and [*ESI*] * *k*_*r*4_, respectively (Figure 7B).

Finally, the product [P] dynamics that are described by equation (12) are also described by the dynamics of voltage [P] in the circuit (Figure 7B). As illustrated above, all the equations (12 and 20-25) describing the reaction kinetics/dynamics for the reaction scheme (Figure 7A) are exactly mapped to a single circuit (Figure 7B). Given the rate constants and initial conditions, these equations (12 and 20-25) can be solved by running simulations of the circuit. In short, this circuit not only accurately visualizes all differential equations in one diagram but can also easily provide solutions to these equations via simulations of the circuit.

Since we haven’t applied any assumptions while deriving the equations or in developing the circuit (Figures 7A, B), this circuit is very general and can be used to model the kinetics of any reaction with the same topology. However, we need to know the values of all the rate constants to run the simulation of this circuit, which is not very convenient. To make the circuit more useful in practice, we simplified the reaction by applying the steady-state approximation (Figure 7C), which is also used in the derivation of enzyme inhibition kinetics from the Michaelis-Menten equation. Under this approximation, substrate binding and inhibitor binding to an enzyme are viewed as instantaneous (much faster than the catalytic reaction) and thus all intermediate complexes reach quasi-steady states. Therefore, we have the equilibrium equations below:

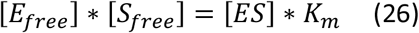

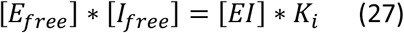

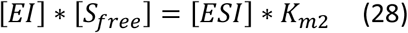

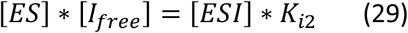

Similar to the tactics used to derive the Michaelis-Menten circuit (Figure 3D), these equilibrium equations are reflected in the circuit (Figure 7D) by removing all capacitors, setting all four *k*_*f*_ values equal to 1, and combining currents for each complex into one or two resistors. We thus achieve a parameter-reduced circuit generalized for enzymatic reactions with inhibition (Figure 7D). Note that ESI has two production fluxes, one from ES and the other from EI, and two consumption fluxes, dissociation from ESI to EI and to ES, respectively. We can sum equations (28) and (29), and have equation (30) which determines the steady state of [*ESI*]:

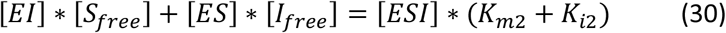

Accordingly, equation (30) is also reflected in the circuit (Figure 7D) for the voltage of [*ESI*], which is determined by two source currents and two sink currents through the two resistors. The two source currents are provided through the *k*_*f*2_ current source and *k*_*f*4_ current source; the two sink currents are realized by the 1/*K*_*m*2_ resistor and 1/*K*_*i*2_ resistor, respectively.

Based on the mathematical derivations above, we have developed a parameter-reduced circuit (Figure 7D) generalized for all four kinds of enzyme inhibition including competitive, non-competitive, uncompetitive, and mixed inhibition. By default, the generalized circuit (Figure 7D) can be viewed as a circuit model for mixed inhibition while the other three types of inhibition are just special cases. For competitive inhibition, since there is only one [*EI*] reaction (Figure 7C), setting the parameters *k*_*f*2_ = *k*_*f*4_ = 0 gives [*ESI*] = 0 and the circuit (Figure 7D) becomes the same circuit in Figure 5B for competitive inhibition. For non-competitive inhibition, the inhibitor binds to the enzyme at a different site from the catalytic site and this binding is independent of the substrate binding, resulting in *K*_*m*2_ = *K*_*m*_ and *K*_*i*2_ = *K*_*i*_ in the reaction scheme (Figure 7C). Consequently, using those same settings in the circuit (Figure 7D), we have the circuit for non-competitive inhibition. Since non-competitive inhibition is a special case of mixed inhibition, when there are no special requirements on *K*_*i*2_ and *K*_*m*2_, the circuit (Figure 7D) by itself is a model for mixed inhibition. For uncompetitive inhibition, there is only one production flux for ESI (from ES), meaning that *k*_*f*3_ = *k*_*f*4_ = 0 in the circuit (Figure 7D). As a result, there is only one consumption flux for [*ESI*] in the uncompetitive inhibition circuit, such that there is only current through the resistor (R = 1/*K*_*i*_), achieved by either removing the other resistor R = 1/*K*_*m*_ or setting its resistance to a huge value such that the current running through it can be ignored. In summary, we have developed a generalized circuit for four types of enzyme inhibition. Its simulation requires five parameters (*K*_*m*_, *k*_*cat*_, *K*_*i*_, *K*_*m*2_ and *K*_*i*2_) for modeling mixed inhibition and only three kinetic parameters, *K*_*m*_, *k*_*cat*_, and *K*_*i*_, for modeling competitive, non-competitive, or uncompetitive inhibition.

The generalized circuit (Figure 7D) consists of two parts: a binding block and a catalytic reaction block. The binding block is the circuit representing the binding of enzyme-substrate (ES), enzyme-inhibitor (EI), and enzyme-substrate-inhibitor (ESI) in Figure 7D. Since the binding circuit can fundamentally simulate different bindings among biomolecules, it is a useful building block for modeling more complicated interactions. Therefore, we extracted the binding block and rearranged the generalized circuit (Figure 7D) into a more concise schematic symbol (Figure 8A) for easy visualization. The new circuit consists of a binding block (blue box) and a catalytic reaction block (Figure 8A), describing exactly the same kinetics/dynamics as in the original circuit (Figure 7D). This binding block has three inputs ([E_0_], [S_0_], and [I_0_]) and three outputs ([ES],[EI], and [ESI]), which are the same as those in Fig 7D. The reaction block is separated; hence there is product [P] feedback to the substrate input to maintain mass conservation of S as shown in Figure 8A. With all the underlying schematic circuits built within it (Figure 7D), our binding-block circuit motif symbol is convenient to use: As in highly complex electronic integrated-circuit design with hierarchical building blocks, it provides a useful visualization aid; it can be repeatedly instantiated to create ever-more complex biochemical reaction networks.

**Figure 8.**
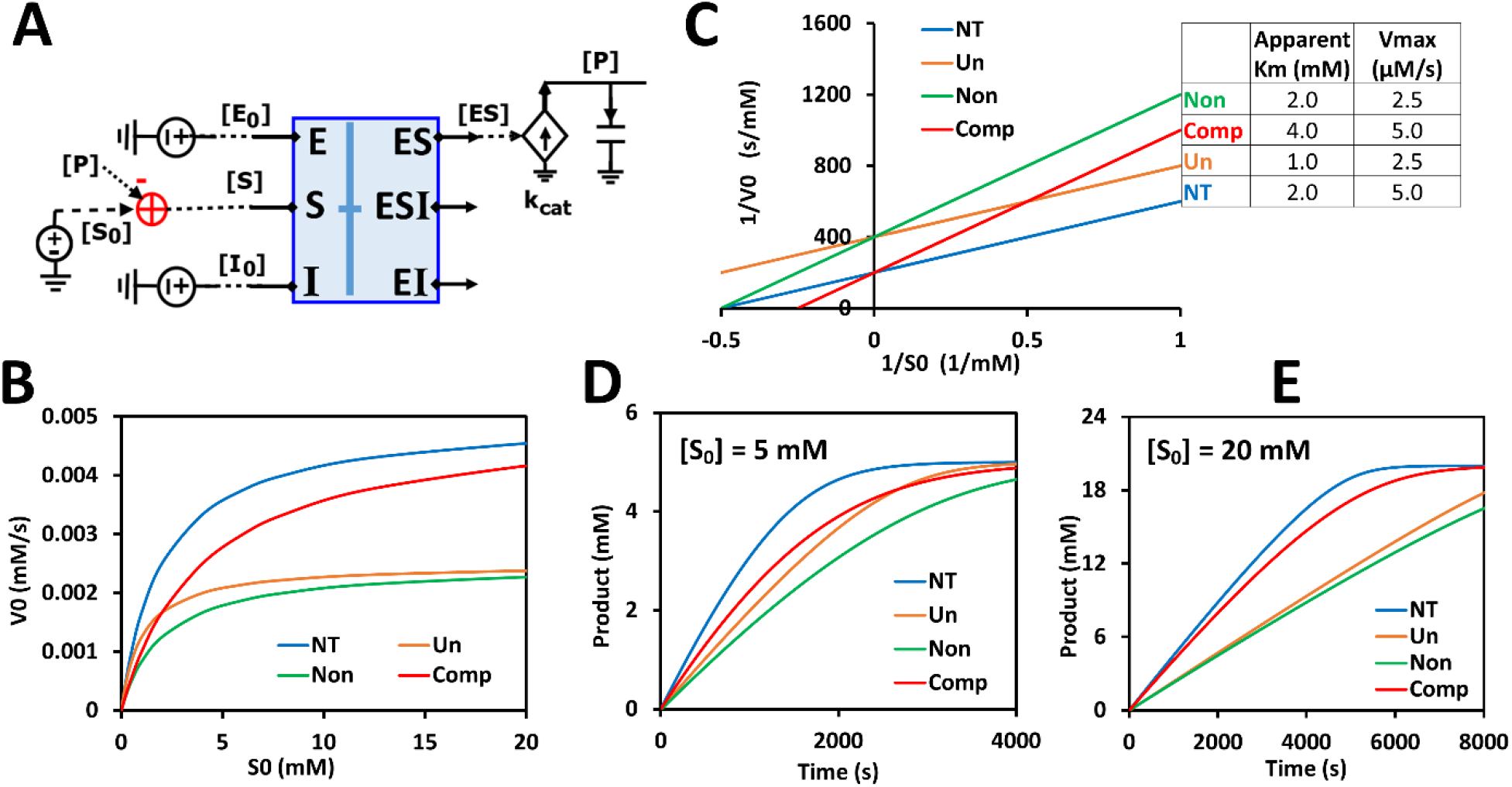
Generalized circuit simulations of enzymatic reactions with different inhibition types. The kinetic parameters used in the simulations are K_m_ = 2 mM, K_i_ = 3 mM, k_cat_ = 500/s, E_0_ = 10 nM, I_0_ = 3 mM, with varying amounts of substrate S. (A) The simplified circuit schematic for the generalized inhibition circuit from Figure 7D with a binding block (blue box) and a catalytic reaction. The binding block is the same circuit used for the binding of enzyme, inhibitor, and substrate in Figure 7D, with a separated catalytic reaction block. (B) Simulation curves of initial substrate concentrations S_0_ versus initial reaction rates V_0_ for enzymatic reaction without inhibitor (NT), with an uncompetitive inhibitor (Un), with a noncompetitive inhibitor (Non), and a competitive inhibitor (Comp). (C) Lineweaver-Burk plot plotting 1/S_0_ versus 1/V_0_. The inserted table corresponds to parameters derived from the X- and Y-intercepts of the corresponding line. These derived parameters exactly match the input parameters. (D,E) Predicted product dynamics over time with S = 5 mM (D) and S = 20 mM (E) for all inhibition types listed.

To verify that our generalized circuit can accurately simulate all enzyme inhibition types, we ran simulations with the circuit using the same kinetic parameters (*K*_*m*_, *k*_*cat*_, and *K*_*i*_) and initial concentrations of enzyme, substrate, and inhibitor for all of its special cases, i.e., for competitive, noncompetitive, and uncompetitive inhibition. As expected, our circuit model predicted the characteristic curves for the relationship between [S_0_] and initial reaction rate V_0_ for all three inhibition types (Figures 8B). Lineweaver-Burk plotting further confirmed that our circuit model is accurate (Figure 8C): For competitive inhibition, the apparent K_m_ increases but V_max_ remains the same upon the addition of the inhibitor; for non-competitive inhibition, the V_max_ decreases but apparent K_m_ remains the same; for uncompetitive inhibition the apparent K_m_ and V_max_ both decrease with the same proportion such that the curve’s slope (K_m_/V_max_) without inhibitor (NT, the blue line in Figure 8C) is identical to that with inhibitor (Un, the orange line in Figure 8C). Furthermore, this circuit can also simulate mixed inhibition where K_m_, K_m2_, K_i_, and K_i2_ are independent of each other, the most general case. As an example, when using the same K_m_ and K_i_ values as in Figure 8 but setting K_i2_ = 6 mM and K_m2_ = 4 mM, we obtain a typical curve for mixed inhibition with different X- and Y-intercepts on the Lineweaver-Burk plot (Figure S1).

Remarkably, this circuit shows that the time-course dynamics of the product differ greatly amongst the three inhibition types (Figures 8D,E) and that the difference depends on substrate concentration: When S = 5 mM, the product dynamics of uncompetitive inhibition are similar to that of competitive inhibition (Figure 8D), while when S = 20 mM the product dynamics of uncompetitive inhibition are closer to that of noncompetitive inhibition (Figure 8E). Non-competitive inhibition usually has the strongest inhibition among the three types, with the slowest product production (Figures 8B,D,E). Our simulation results suggest that, to obtain accurate molecular kinetics, it is important to choose the right inhibition mechanism when constructing models. Choosing the right type of inhibition is especially critical for cascade reactions such as in metabolic pathways where there are many steps with feedback inhibition caused by the final product and/or intermediate metabolites.

### 2.6 Circuit modeling of non-competitive inhibition

We have now verified in theory that the generalized circuit (Figure 8A) is accurate in predicting kinetics for all enzyme inhibition types. This circuit is also convenient to use; with inputs of K_m_, k_cat_ and K_i_ as well as experimental initial conditions, running simple simulations can provide all kinetic data, which can be compared to experimentally measured data. As an example, we now use our circuit to model the noncompetitive inhibition wherein lactate oxidation by lactate dehydrogenase is inhibited by a noncompetitive inhibitor, oxamate (Powers et al., 2007). In this redox reaction (Figure 9A), we varied the concentration of lactate while keeping a constant NAD^+^ concentration. As expected, the circuit-predicted curves capture the experimental data nicely. The initial reaction rate increases as substrate concentration increases, but, in the presence of the noncompetitive inhibitor, the reaction rate never approaches the V_max_ of the reaction when without the inhibitor (Figure 9B). Linearized curves from Lineweaver-Burk plots show that noncompetitive inhibition has the same K_m_ as the reaction without the inhibitor, but a smaller V_max_ (i.e., greater 1/V_max_) (Figure 9C). The generalized circuit can also simulate the time-course kinetics of NADH production for the initial period (the first 360s) (Figure 9D) where the catalytic reaction can be largely viewed in the forward direction.

**Figure 9.**
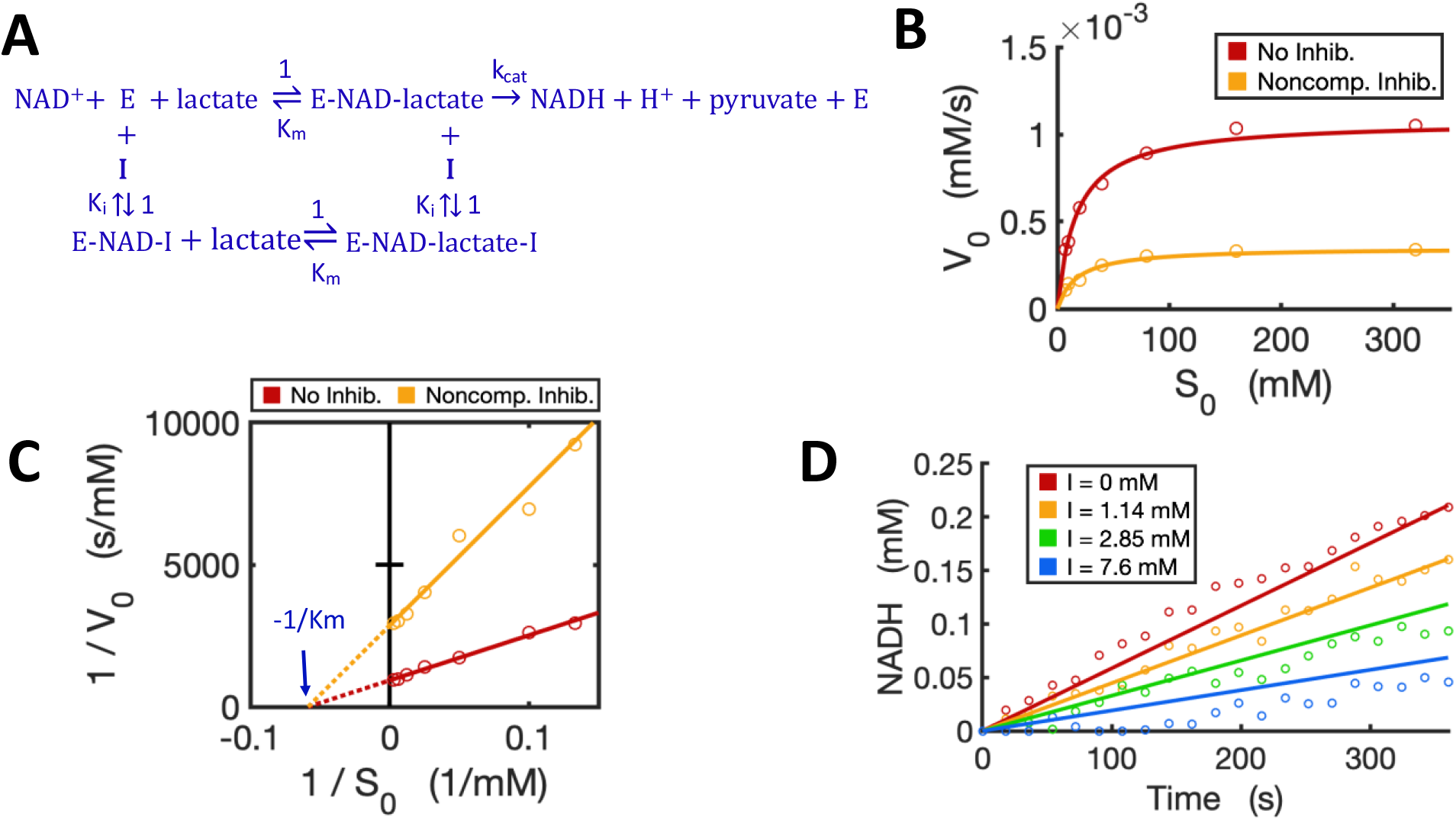
Circuit modeling of non-competitive inhibition of LDH by oxamate. (A) The reaction scheme of non-competitive inhibition against LDH, where the inhibitor I is oxamate. For simplification, the initial catalysis can be viewed as a directional reaction, though the LDH reaction is known to be reversible; the fast binding step of NAD^+^ to enzyme is neglected in this reaction scheme. (B) Simulation curves of V_0_ versus S_0_ fitted to measured experimental data in the absence and presence of oxamate (7.6 mM), when E_0_ = 5.0 nM. (C) Lineweaver-Burk plot showing linearized curves of 1/V_0_ versus 1/S_0_ with and without the inhibitor, which have the same X-intercept of - 1/K_m_. (D) The model curves of product dynamics over time with varying inhibitor concentrations fitted to experimental data points (when S = 80 mM, E_0_ = 3.3 nM). All simulation curves are obtained from the circuit model (Figures 7 and 8A) with experimentally measured K_m_ = 17.1 mM, k_cat_ = 215.5 (1/s), and K_i_ = 3.66 mM. All data points are means of three independent replicates with standard deviation less than 20% of the corresponding mean (not shown).

### 2.7 Circuit modeling of reversible reactions

We developed a circuit to simulate reversible reactions because they are very common in metabolic pathways. Since the kinetic mechanism for the reversible reaction of alcohol dehydrogenase (ADH) is well studied with all rate constants known (Dickinson and Dickenson, 1978; Ganzhorn et al., 1987; Plapp, 2010), we developed a circuit model for this reversible reaction with ethanol and NAD^+^ both as substrates. The kinetics of this reaction by yeast ADH follow the ordered Bi-Bi mechanism, resulting in a five-step reversible reaction (Ganzhorn et al., 1987) as shown in Figure 10A. First, NAD+ binds to the enzyme ADH and forms an E-NAD complex; the substrate ethanol then binds to E-NAD and forms an E-NAD-S complex; this complex turns into a new complex E-NADH-P and then the first product aldehyde (P) releases, resulting in an E-NADH complex; the last step is the dissociation of NADH from the enzyme. The total reaction consists of five reversible steps, each with forward and reverse rate constants as indicated in the reaction scheme (Figure 10A).

**Figure 10.**
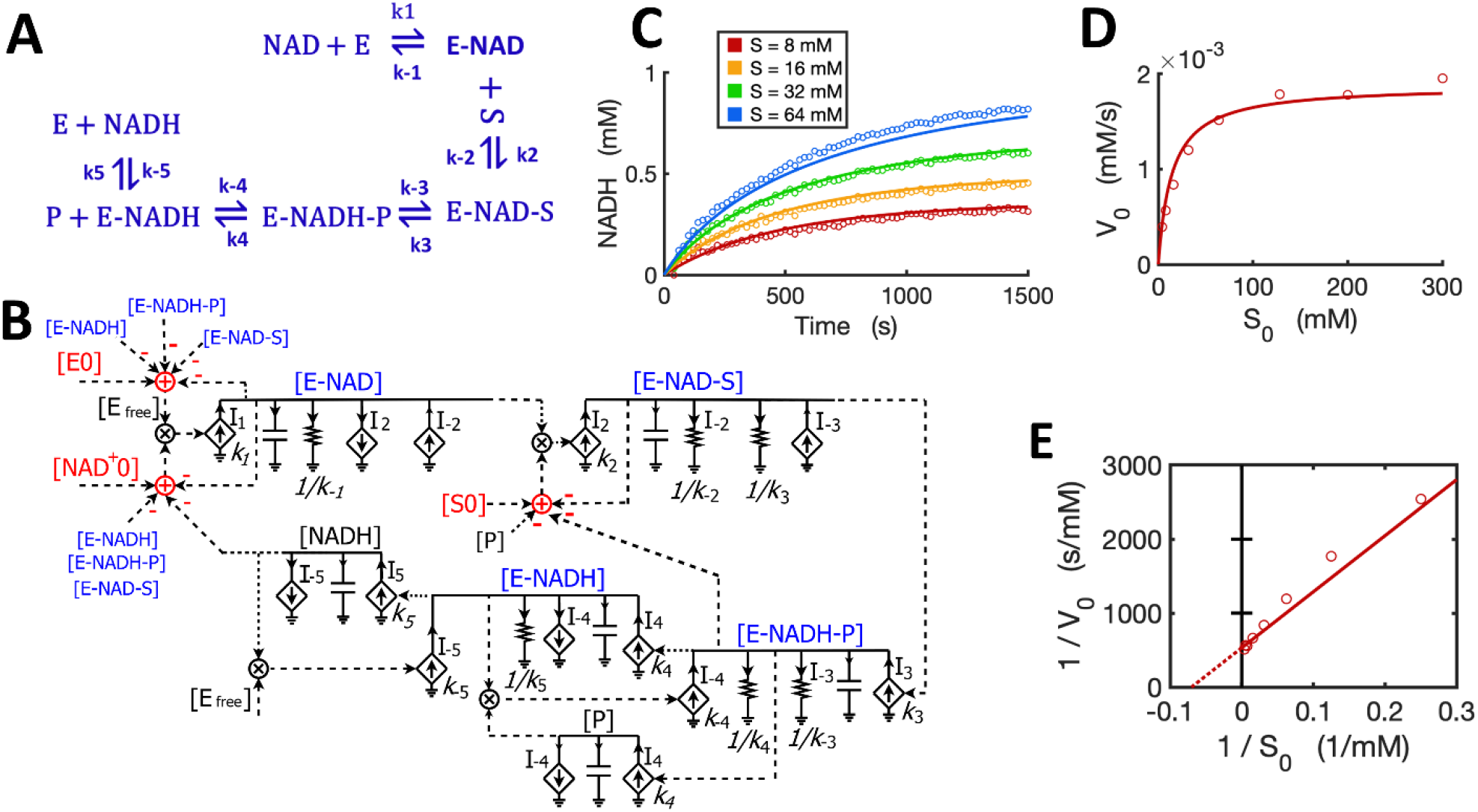
Circuit modeling of a reversible reaction catalyzed by yeast alcohol dehydrogenase (ADH). (A) The mechanistic scheme for a five-step reversible reaction for yeast ADH, where E is ADH enzyme, S is ethanol, and P is the acetaldehyde produced. (B) Circuit model that exactly matches the kinetics of the reaction scheme in (A). [E_0_] is the initial ADH concentration (3.9 nM), [NAD^+^_0_] is the initial NAD^+^ concentration (4 mM), [S_0_] is the initial ethanol concentration. Arrows along solid lines or in current generators (diamond symbols) indicate the direction of the corresponding current (reaction flux). Voltages or currents labeled with the same names indicate the same values. All circuit symbols are shown in Figures 1 and 3. (C) Model curves of product [NADH] dynamics over time fitted to experimental data points with varying ethanol concentrations. (D) Simulation curves of initial reaction rates V_0_ versus initial alcohol concentrations S_0_ fitted to experimental data. (E) Lineweaver-Burk plot showing a straight curve of 1/V_0_ versus 1/S_0_. All data points are means of three independent replicates with standard deviations less than 20% of the corresponding mean (not shown).

To mathematically simulate the dynamics/kinetics of all species in this multi-step reaction would require a long list of differential equations as well as many mass conservation equations. Instead, we can simply translate the reaction scheme into a single circuit (Figure 10B) without deriving differential equations. The circuit model is constructed by using a circuit block similar to the one for the enzyme-substrate circuit (Figure 3B). The difference is that each intermediate complex (blue text) including E-NAD, E-NAD-S, E-NADH-P, and E-NADH in steps 1-4 has two production fluxes and two consumption fluxes (Figure 10B), instead of only three fluxes for the [ES] complex in the circuit (Figure 3B) for classic Michaelis-Menten kinetics without a reversible catalytic reaction. It should be noted that the product aldehyde [P] is produced in proportion to [E-NADH-P] with a rate constant of k_4_ while it is consumed by binding back to [E-NADH], forming [E-NADH-P] with a rate constant k_-4_, resulting in a reaction rate of I_-4_ = k_-4_*[P]*[E-NADH]. Similarly, another product [NADH] is produced in proportional to [E-NADH] with a reaction rate of k_5_*[E-NAD] while it is also consumed by binding back to the enzyme, forming [E-NADH] with a rate of I_-5_ = k_-5_*[NADH]*[E_free_]. Another important note is that the conservations of enzyme, NAD+, and ethanol (S) during the reaction are indicated by adder and subtraction symbols in red in the circuit (Figure 10B).

To verify the accuracy of our circuit model, we fit experimental data for [NADH] produced from ethanol oxidation by ADH for varying amounts of alcohol and a fixed concentration of initial NAD^+^. Given that the 10 rate constants for the whole reaction are known from previous studies (Dickinson and Dickenson, 1978; Ganzhorn et al., 1987), we used these kinetic parameters as input to our circuit model. We were able to get a good fit with only some adjustments (Table S1 in the Supplementary) relevant to our specific experimental conditions. Our circuit model can accurately simulate the product dynamics over time with varying amounts of ethanol input (Figure 10C). Importantly, our circuit model also correctly predicts the classic relationship between initial reaction rates (V_0_) and initial ethanol concentrations (S_0_) based on modeling the initial 60 s of the reaction (Figures 10D). The Lineweaver-Burk plot shows a straight line for 1/V_0_ versus 1/S_0_, which captures our data closely (Figure 10E). From the X- and Y-intercepts, we calculated a K_m_ of 14.3 mM for ethanol and k_cat_ of 483/s, which agrees well with previous reports (Ganzhorn et al., 1987).

### 2.8 Circuit modeling of transcription and translation in a cell-free system

Circuits can be used to model any biological system, not just enzymatic reactions (Sarpeshkar, 2010; Teo and Sarpeshkar, 2020). Here, we sought to model transcription and translation (TXTL) kinetics regulated by TetR in an *E. coli*-based cell-free system. In this system, the DNA insert region (a hybrid T7 promoter regime) on a plasmid (DA313) has a T7 RNAP binding site and a TetR binding site (tetO) that is 5-bp downstream; given the relatively small sizes of T7RNAP and TetR, the DNA region acts like an enzyme capable of binding both the “substrate” (T7 RNAP) and the inhibitor protein (TetR) at two different sites. Since T7 RNAP and the inhibitor TetR bind to the DNA regime (“enzyme”) at two different sites, we can assume their interactions to architect noncompetitive binding (Figure 11A). Such transcriptional binding is similar to the noncompetitive binding/reaction scheme (Figures 7C,D and 8A) but with the difference that none of the “substrate” (T7 RNAP) is consumed or converted into products. The free DNA-RNAP complex turns on the transcription followed by the translation of GFP (Figure 11 A). The dynamics of mRNA are determined by production and degradation, with rate constants of k_TX_ (1/s) and d (1/s), respectively. Ribosomes then bind to the ribosome binding site (RBS) of the mRNA and turn on the translation of GFP_dark with a rate constant of k_TL_ (1/s). GFP_dark undergoes folding and maturation with a rate constant of k_mat_ (1/s), resulting in fluorescent GFP (Figure 11 A). In this study, we focused only on the initial TXTL reactions in the first ∼4000 s, wherein the reactions reach their maximal production rates (steady states) and are not limited by the available amino acids, energy sources, and/or other components. Since TetR forms stable homodimers (Krafft et al., 1998), TetR in this model represents its homodimer. With these simplifications and assumptions, we thus have the reaction scheme for TXTL regulated by TetR in the cell-free system (Figure 11A).

**Figure 11.**
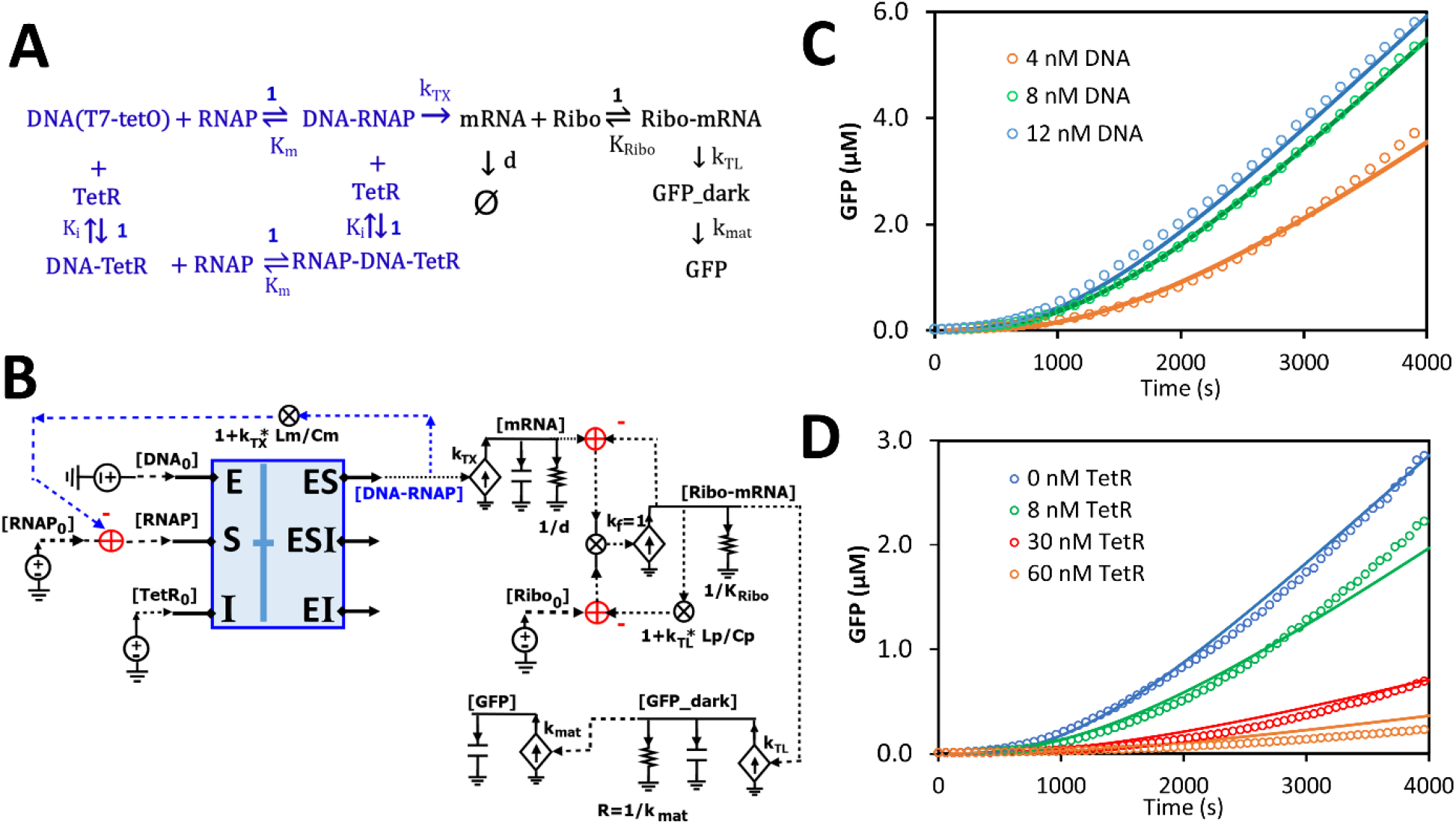
Circuit modeling of cell-free transcription and translation regulated by TetR. (A) The scheme for molecular binding and associated reactions of TXTL in the cell-free system with repression by TetR. The TXTL is turned on by a hybrid T7 promoter (DNA) and T7 RNA polymerase (RNAP) and repressed by TetR via non-competitive inhibition. The free RNAP-DNA complexes turn on transcription followed by the translation of GFP. The TetR used in the model is its homodimer. K_i_ represents the dissociation constant (K_d_) of the homodimer binding to the DNA (tetO). (B) The circuit model exactly describes the kinetics of molecular binding and associated reactions in (A). In the non-competitive binding block (blue box), DNA acts in a manner like an enzyme where both the substrate (RNAP) and the inhibitor (TetR) bind to it. Only free [DNA-RNAP] turns on transcription. (C) Model curves of GFP dynamics fitted to experimental data points with varying DNA concentrations when TetR = 0 nM. (D) Model curves of GFP dynamics fitted to experimental data points with varying TetR concentrations at constant DNA concentration (6 nM). All data points are means of three independent replicates with standard deviations less than 20% of the corresponding mean (not shown).

Based on the binding and reaction scheme (Figure 11A), we used the general binding block (Figure 8A) and designed a circuit (Figure 11B) that exactly describes the kinetics of the binding interactions and reactions for the regulated TXTL in the cell-free system. In this circuit (Figure 11B), the binding block (blue box) has voltage inputs of initial [DNA_0_] for total plasmid concentration, [RNAP_0_] for total T7 RANP concentration, and [TetR_0_] for total TetR concentration, via the pins of E, S, and I, respectively; free [TetR] and [RNAP] bind to DNA noncompetitively, exactly described by the same circuit as Figure 7C or 8A; among the three outputs, only the [DNA-RNAP] turns on the transcription; the mRNA is then bound by ribosomes, which results in GFP production. Since several RNAP molecules can bind to one DNA molecule during transcriptional elongation, the average number of T7-RNAP bound in each DNA-RNAP complex would be 1+ k_TX_*Lm/Cm where 1 is the one RNAP binding to the promoter, K_TX_ is the transcription rate (1/s), Lm is the length of mRNA (nt), and Cm is the average RNAP velocity during elongation (nt/s) (Bremer et al., 2003; Marshall and Noireaux, 2019). Therefore, for each DNA-RNAP complex the number of RNAP, 1+ k_TX_*Lm/Cm, has to be subtracted from the total [RNAP_0_] pool. Similar to the counting of RNAP copy number on each DNA-RNAP complex, for each Ribo-mRNA complex, there are 1+ k_TX_*Lp/Cp ribosomes, where 1 is the one ribosome binding to the ribosome binding site, k_TL_ is the translation rate (1/s), Lp is the GFP coding length of mRNA (nt), and Cp is the ribosome velocity during elongation (nt/s). The last part of the circuit is the GFP production where GFP_dark is produced with a rate constant of k_TL_ and consumed to make fluorescent GFP with a rate constant of k_mat_.

All the kinetic parameters (Table S2) used in this model were obtained either from previous studies (Újvári and Martin, 1996; Kamionka et al., 2004; Skinner et al., 2004; Marshall and Noireaux, 2019) under similar conditions or from fitting to our experimental data. The simulation curves fitted our experimental data closely as shown in Figures 11C and 11D. As expected, as more plasmid was added to the cell-free system more GFP was produced until saturation was reached (up to ∼12 nM DNA in our experiments) (Figure 11C), where adding more plasmid would not increase GFP production. In another experiment, when varying amounts of TetR were added to the cell-free system with a constant DNA concentration, GFP production was increasingly repressed by more TetR addition.

## Discussion

We have shown that the ordinary differential equations used to describe the dynamics of voltages and currents in relatively simple electronic circuits are exactly the same as the ODEs describing molecular kinetics in chemical reactions and biological systems. Voltages faithfully represent molecular concentrations while currents faithfully represent reaction fluxes. Furthermore, we have built Michaelis-Menten circuits and validated them by using well-defined parameters and by fitting experimental data from enzymatic reactions. Based on the MM circuits, we then developed circuit models for more complicated reactions including various enzyme inhibition types, product feedback inhibition, reversible reactions, and regulated TXTL in a cell-free system. These circuits can provide foundational building blocks for kinetic modeling of complex chemical and biological networks.

Taking advantage of circuit models, we are able to translate reaction schematics directly into circuits and to directly perform rapid kinetic modeling for biochemical reactions/networks without the need for math equations. Using electronic circuits as a new modeling language can thus enable faster and easier modeling of molecular kinetics than using ODEs and numerical solvers. Importantly, circuit modeling is also very effective for scientific communication. Circuit modeling is not only accurate but also very concise because all the underlying equations and parameters/terms describing molecular kinetics are visualized and clearly labeled in one circuit.

It should also be noted that our circuit models are more general and accurate than classic MM kinetics because we don’t need the free ligand assumption where [S_0_] is assumed to be much greater than [E_0_]. Instead, subtraction of [ES] and [P] from [S_0_] in all of our circuits always ensures exact mass conservation. Similarly, if present, automatic subtraction of [EI] and [ESI] ensures exact mass conservation. These subtractions may not necessarily improve the model accuracy for normal enzymatic reactions that are run under conditions of [S_0_] >> [E_0_] and/or [I_0_] >>[E_0_]. However, under some circumstances where the concentrations of substrate and/or inhibitor are close to the enzyme concentration, such as the case of our TXTL modeling in the cell-free system (Figure 10), the concentrations of complexes including [ES], [EI] and/or [ESI] account for considerable portions of the total concentrations of [S_0_], [I_0_] and/or [E_0_]. Thus, with built-in accounting for the concentrations of these complexes, we can apply circuits more accurately and conveniently than MM equations that may not be accurate under all conditions.

## Conclusion

Circuit modeling is advantageous for rapid and accurate kinetic modeling of chemical and biological systems. Circuit models visualize all math equations and their relationships in one circuit, which is very concise and convenient for effective communication. We envision that circuit modeling could be adopted as a new language for kinetic modeling and promote scientific communication in the fields of biochemistry, systems biology, and synthetic biology.

## Methods and Materials

All enzymes and chemicals were purchased from Sigma Aldrich unless otherwise mentioned. Three enzymes used in this study were beta-galactosidase from *E. coli* (cat# G5635), rabbit muscle lactate dehydrogenase (cat# L2500), and yeast alcohol dehydrogenase (cat# A7011).

### Enzymatic assays

All enzymatic assays were run in 96-well plates (Costar, cat#3595) with 100 μl of reaction mixture per well and reactions were monitored in real time by using a microplate reader (Molecular Devices Inc.). Beta-galactosidase (beta-Gal) assays were performed using o-nitrophenyl-β-D-galactopyranoside (ONPG) as substrate. All reagents were prepared in PBS buffer (pH7.2) with 5 mM Dithiothreitol (DTT) and 4 mM MgCl_2_. The reactions (100 μl/well) with varying concentrations of ONPG were monitored by measuring the absorbance of the product, 2-nitrophenol, at 420 nm in real time for over 20 minutes. The concentrations of the product were determined by a calibration curve that was made by using a freshly prepared 2-nitrophenol solution under the same condition. Initial reaction rates were calculated from the slopes of each reaction curve normally within the first 6-8 minutes. For competitive inhibition, phenylethyl beta-D-thiogalactopyranoside (PETG), a known competitive inhibitor of beta-Gal (Xu and Ewing, 2004), was added to the reactions under the same condition. For the product feedback inhibition, galactose at varying concentrations was added to the reaction mixture before running the enzymatic assay.

The lactate dehydrogenase (LDH) assay was performed in 96-well plates using a constant concentration of 2mM NAD+ and varying the concentration of lactate as substrate. All reagents were prepared in 0.5 M glycine buffer (pH9.5). The end product NADH concentration was determined by measuring absorbance at 350 nm with a calibration curve pre-established under the same condition. All the reactions (100 μl/well) were monitored in real time in 96-well plates by a microplate reader. For modeling non-competitive inhibition, oxamate was used as a non-competitive inhibitor of LDH, in the lactate oxidation direction (Powers et al., 2007). The alcohol dehydrogenase (ADH) assay was performed in 40 mM Tris buffer (pH8.3) by using a constant concentration of 4 mM NAD+ and varying the concentration of ethanol as substrate. The reactions were monitored similarly to the LDH assay. All experiments were done with three replicates unless otherwise mentioned. Control reactions without any enzyme were also included in each experiment as blanks. The kinetic parameters including K_m_, K_i_, V_max_, and k_cat_, were derived from Lineweaver-Burk plotting. These experimentally obtained kinetic parameters and initial concentrations were put into the corresponding circuit models before running simulations.

### Transcription and Translation in the cell-free system

The cell-free *E. coli* protein synthesis system was purchased from New England BioLabs (cat# E5360). The T7 RNA Polymerase (NEB, cat# M0251S) and RNase Inhibitor (NEB, cat#M0314S) were added to the reaction mixture. The transcription and translation reactions (15 μl/each) with varying concentrations of plasmid DA313 and purified TetR were run in a 384-well plate (ThermoFisher Scientific, cat# 12-566-2). The reactions were monitored in real time by measuring GFP fluorescence at Ex485/Em528 in a microplate reader (Molecular Devices Inc.). The plasmid DA313 was made by fusing fragments of a hybrid T7 promoter with a tetO binding site (Jung et al., 2020), 5’UTR sequence, and a superfolder GFP (sfGFP) gene into the backbone of pJBL7010 (Silverman et al., 2019) by Gibson assembly (NEB, cat# E2621L). The 5’UTR sequence includes an mRNA stability hairpin (Carrier and Keasling, 1999; Silverman et al., 2019) and a ribosome binding site that was designed by the RBS calculator (Salis, 2011). The plasmid DA313 was purified by Monarch plasmid miniprep kit (New England BioLabs, cat# T1010S) and its concentration was determined by a fluorometric assay with a DNA-specific dye EvaGreen (Ihrig et al., 2006). The sfGFP concentration was estimated by a pre-established calibration curve using a pure EGFP (BioVision, cat#4999-100) with a conversion factor of 1.6 to account for different fluorescent brightness between sfGFP and EGFP (Fluorescent Protein Database, https://www.fpbase.org/). All experiments were performed with three independent replicates.

The recombinant TetR with 6xHis tag was over-expressed in *E. coli* JM109 (DE3) with plasmid DA303 and purified using Nickel-NTA agarose resin (ThermoFisher Scientific, cat# 88221). The purity of the recombinant TetR was analyzed by SDS-PAGE gel and the concentration was measured by BCA assay (ThermoFisher Scientific, cat#23252). Since TetR forms stable homodimers (Krafft et al., 1998) its concentration is then converted to the molar concentration of the homodimer. The plasmid DA303 was made by fusing TetR into the pQE80 backbone by Gibson assembly. All plasmids used were confirmed by Sanger sequencing and their maps are provided in the supplemental material (Figure S2).

## Data Availability Statement

All supporting data generated or analyzed in this study are included in this published article and the supplementary material or available upon request to the corresponding author.

## Author Contributions

Y.D., D.R.B., and R.S. conceived and designed the circuits and projects; Y.D. and X.R. conducted the experiments and collected all the data; Y.D. and D.R.B analyzed the data, and performed circuit modeling and simulation; Y.D. and T.G.R. made the circuit schematics; Y.D. wrote the manuscript and all authors revised and approved the final manuscript.

## Supplementary Material

The Supplementary Material for this article can be found online at:

## Publisher’s Note

All claims expressed in this article are solely those of the authors and do not necessarily represent those of their affiliated organizations, or those of the publisher, the editors and the reviewers. Any product that may be evaluated in this article, or claim that may be made by its manufacturer, is not guaranteed or endorsed by the publisher.

## References

Alves, R., Antunes, F., and Salvador, A. (2006). Tools for kinetic modeling of biochemical networks. Nat. Biotechnol. 24, 667–672. doi:10.1038/nbt0606-667.

Beahm, D. R., Deng, Y., Riley, T. G., and Sarpeshkar, R. (2021). Cytomorphic Electronic Systems: A review and perspective. IEEE Nanotechnol. Mag. 15, 41–53. doi:10.1109/MNANO.2021.3113192.

Bevc, S., Konc, J., Stojan, J., Hodošcek, M., Penca, M., Praprotnik, M., et al. (2011). ENZO: A web tool for derivation and evaluation of kinetic models of enzyme catalyzed reactions. PLoS One 6. doi:10.1371/journal.pone.0022265.

Bremer, H., Dennis, P., and Ehrenberg, M. (2003). Free RNA polymerase and modeling global transcription in Escherichia coli. Biochimie 85, 597–609. doi:10.1016/S0300-9084(03)00105-6.

Carrier, T. A., and Keasling, J. D. (1999). Library of synthetic 5’ secondary structures to manipulate mRNA stability in Escherichia coli. Biotechnol. Prog. 15, 58–64. doi:10.1021/bp9801143.

Daniel, R., Rubens, J. R., Sarpeshkar, R., and Lu, T. K. (2013). Synthetic analog computation in living cells. Nature 497, 619–623. doi:10.1038/nature12148.

Deng, Y., Beahm, D. R., Ionov, S., and Sarpeshkar, R. (2021). Measuring and modeling energy and power consumption in living microbial cells with a synthetic ATP reporter. BMC Biol. 19, 1–21. doi:10.1186/s12915-021-01023-2.

Dickinson, F. M., and Dickenson, C. J. (1978). Estimation of rate and dissociation constants involving ternary complexes in reactions catalysed by yeast alcohol dehydrogenase. Biochem. J. 171, 629–37. doi:10.1042/bj1710629.

Eshtewy, N. A., and Scholz, L. (2020). Model reduction for kinetic models of biological systems. Symmetry (Basel). 12, 1–22. doi:10.3390/SYM12050863.

Ganzhorn, A. J., Green, D. W., Hershey, A. D., Gould, R. M., and Plapp, B. V. (1987). Kinetic characterization of yeast alcohol dehydrogenases. Amino acid residue 294 and substrate specificity. J. Biol. Chem. 262, 3754–3761. doi:10.1016/s0021-9258(18)61419-x.

Ihrig, J., Lill, R., and Mühlenhoff, U. (2006). Application of the DNA-specific dye EvaGreen for the routine quantification of DNA in microplates. Anal. Biochem. 359, 265–267. doi:10.1016/j.ab.2006.07.043.

Jung, J. K., Alam, K. K., Verosloff, M. S., Capdevila, D. A., Desmau, M., Clauer, P. R., et al. (2020). Cell-free biosensors for rapid detection of water contaminants. Nat. Biotechnol. 38, 1451– 1459. doi:10.1038/s41587-020-0571-7.

Kamionka, A., Bogdanska-Urbaniak, J., Scholz, O., and Hillen, W. (2004). Two mutations in the tetracycline repressor change the inducer anhydrotetracycline to a corepressor. Nucleic Acids Res. 32, 842–847. doi:10.1093/nar/gkh200.

Krafft, C., Hinrichs, W., Orth, P., Saenger, W., and Welfle, H. (1998). Interaction of Tet repressor with operator DNA and with tetracycline studied by infrared and Raman spectroscopy. Biophys. J. 74, 63–71. doi:10.1016/S0006-3495(98)77767-7.

Marshall, R., and Noireaux, V. (2019). Quantitative modeling of transcription and translation of an all-E. coli cell-free system. Sci. Rep. 9, 11980. doi:10.1038/s41598-019-48468-8.

Néant, N., Lingas, G., Le Hingrat, Q., Ghosn, J., Engelmann, I., Lepiller, Q., et al. (2021). Modeling SARS-CoV-2 viral kinetics and association with mortality in hospitalized patients from the French COVID cohort. Proc. Natl. Acad. Sci. U. S. A. 118. doi:10.1073/pnas.2017962118.

Nguyen, T.-H., Splechtna, B., Steinböck, M., Kneifel, W., Lettner, H. P., Kulbe, K. D., et al. (2006). Purification and characterization of two novel beta-galactosidases from Lactobacillus reuteri. J. Agric. Food Chem. 54, 4989–98. doi:10.1021/jf053126u.

Plapp, B. V. (2010). Conformational changes and catalysis by alcohol dehydrogenase. Arch. Biochem. Biophys. 493, 3–12. doi:10.1016/j.abb.2009.07.001.

Portaccio, M., Stellato, S., Rossi, S., Bencivenga, U., Mohy Eldin, M. S., Gaeta, F. S., et al. (1998). Galactose competitive inhibition of β-galactosidase (Aspergillus oryzae) immobilized on chitosan and nylon supports. Enzyme Microb. Technol. 23, 101–106. doi:10.1016/S0141-0229(98)00018-0.

Powers, J. L., Kiesman, N. E., Tran, C. M., Brown, J. H., and Bevilacqua, V. L. H. (2007). Lactate dehydrogenase kinetics and inhibition using a microplate reader. Biochem. Mol. Biol. Educ. 35, 287–292. doi:10.1002/bmb.74.

Resat, H., Petzold, L., and Pettigrew, M. F. (2009). “Kinetic Modeling of Biological Systems BT - Computational Systems Biology,” in, eds. R. Ireton, K. Montgomery, R. Bumgarner, R. Samudrala, and J. McDermott (Totowa, NJ: Humana Press), 311–335. doi:10.1007/978-1-59745-243-4_14.

Salis, H. M. (2011). The ribosome binding site calculator. Methods Enzymol. 498, 19–42. doi:10.1016/B978-0-12-385120-8.00002-4.

Sarpeshkar, R. (2010). Ultra Low Power Bioelectronics: Fundamentals, Biomedical Applications, and Bio-Inspired Systems. Cambridge: Cambridge University Press doi:10.1017/CBO9780511841446.

Silverman, A. D., Kelley-Loughnane, N., Lucks, J. B., and Jewett, M. C. (2019). Deconstructing Cell-Free Extract Preparation for in Vitro Activation of Transcriptional Genetic Circuitry. ACS Synth. Biol. 8, 403–414. doi:10.1021/acssynbio.8b00430.

Skinner, G. M., Baumann, C. G., Quinn, D. M., Molloy, J. E., and Hoggett, J. G. (2004). Promoter binding, initiation, and elongation by bacteriophage T7 RNA polymerase: A single-molecule view of the transcription cycle. J. Biol. Chem. 279, 3239–3244. doi:10.1074/jbc.M310471200.

Teo, J. J. Y., Kim, J., Woo, S. S., and Sarpeshkar, R. (2019a). Bio-molecular Circuit Design with Electronic Circuit Software and Cytomorphic Chips. in 2019 IEEE Biomedical Circuits and Systems Conference (BioCAS) (IEEE), 1–4. doi:10.1109/BIOCAS.2019.8918684.

Teo, J. J. Y., and Sarpeshkar, R. (2020). The Merging of Biological and Electronic Circuits. iScience 23, 101688. doi:10.1016/j.isci.2020.101688.

Teo, J. J. Y., Weiss, R., and Sarpeshkar, R. (2019b). An Artificial Tissue Homeostasis Circuit Designed via Analog Circuit Techniques. IEEE Trans. Biomed. Circuits Syst. 13, 540–553. doi:10.1109/TBCAS.2019.2907074.

Teo, J. J. Y., Woo, S. S., and Sarpeshkar, R. (2015). Synthetic Biology: A Unifying View and Review Using Analog Circuits. IEEE Trans. Biomed. Circuits Syst. 9, 453–474. doi:10.1109/TBCAS.2015.2461446.

Újvári, A., and Martin, C. T. (1996). Thermodynamic and kinetic measurements of promoter binding by T7 RNA polymerase. Biochemistry 35, 14574–14582. doi:10.1021/bi961165g.

Xu, H., and Ewing, A. G. (2004). A rapid enzyme assay for β-galactosidase using optically gated sample introduction on a microfabricated chip. Anal. Bioanal. Chem. 378, 1710–1715. doi:10.1007/s00216-003-2317-z.

Zeng, J., Teo, J., Banerjee, A., Chapman, T. W., Kim, J., and Sarpeshkar, R. (2018). A Synthetic Microbial Operational Amplifier. ACS Synth. Biol. 7, 2007–2013. doi:10.1021/acssynbio.8b00138.

