## Supplementary Material for "Rapid modeling of experimental molecular kinetics with simple electronic circuits instead of with complex differential equations"

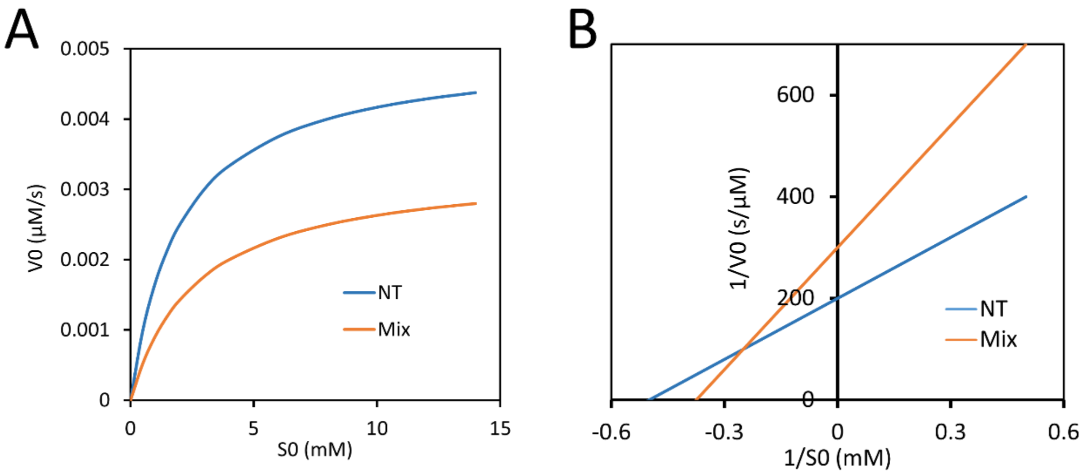


**Figure S1.** Circuit modeling of enzyme inhibition by a mixed-type inhibitor. (A) Model-predicted curves for initial reaction rate ($V_0$) versus initial substrate concentration ($S_0$) with and without inhibitor. (B) Lineweaver-Burk plot for $1/V_0$ versus $1/S_0$ curves, with and without inhibitor, predicted by the circuit model. The model curves are predicted by the circuit in Fig 7D. For comparison, all parameters used in Fig. 8 were kept the same in this figure, including $K_m$ = 2 mM, $K_i$ = 3 mM, $K_{cat}$ = 500/s, $E_0$ = 10 nM, and inhibitor $I_0$ = 3 mM. The extra parameters used for the mixed-inhibition, $K_{i2}$ = 6 mM and $K_{m2}$ = 4 mM, are different from $K_i$ and $K_m$, respectively.

**Table S1. Rate constants used in modeling the reversible reaction by yeast ADH.**

| Rate constants | Values | Cited values | Units | References |
|---|---|---|---|---|
| $k_1$ | $2\text{x}10^6$ | $2\text{x}10^6$ | $M^{-1}s^{-1}$ | (1, 2) |
| $k_{-1}$ | $1.9\text{x}10^3$ | $1.9\text{x}10^3$ | $s^{-1}$ | |
| $k_2$ | $59\text{x}10^3$ | $59\text{x}10^6$ | $M^{-1}s^{-1}$ | |
| $k_{-2}$ | $1\text{x}10^3$ | $1\text{x}10^3$ | $s^{-1}$ | |
| $k_3$ | $8\text{x}10^3$ | $4\text{x}10^3$ | $s^{-1}$ | |
| $k_{-3}$ | $35\text{x}10^3$ | $35\text{x}10^3$ | $s^{-1}$ | |
| $k_4$ | $22\text{x}10^3$ | $11\text{x}10^3$ | $s^{-1}$ | |
| $k_{-4}$ | $4.3\text{x}10^6$ | $4.3\text{x}10^6$ | $M^{-1}s^{-1}$ | |
| $k_5$ | 960 | 480 | $s^{-1}$ | |
| $k_{-5}$ | $15\text{x}10^6$ | $15\text{x}10^6$ | $M^{-1}s^{-1}$ | |

Note: All parameters used in our model are identical or near to values cited in our references except $k_2$, which needed a 1000-fold decrease to give us a good fit to measured experimental data and to yield a $K_m$ for ethanol binding, which is consistent with other measured values (2). The $k_2$ reported in (1), likely an overestimate, leads to a smaller than usual $K_m$ for ethanol binding and saturated the reaction at all ethanol concentrations used in our study.

**Table S2 Kinetic parameters used for TXTL in the *E.coli*-based cell-free system**

| Parameters | | Values used [a, b] | Parameters in circuits [a, b] | Reference values | References |
|---|---|---|---|---|---|
| $K_m$_T7 | Dissociation constant for T7 RNAP and the T7 promoter | 4.8 nM | R = $1/K_m$ = $2.08*10^8$ Ω | 4.8 nM | (3) |
| [$RNAP_0$] | T7 RNAP concentration used | 16 nM | 16 nV | This study | |
| Lm | Total mRNA length | 900 nt | 900 nt | This study (DA313 plasmid) | |
| Lp | GFP coding region length | 717 nt | 717 nt | | |
| Cm | T7 RNAP speed | 50 nt/s | 50 nt/s | 43 nt/s | (4) |
| $k_{TX}$ | Transcription rate | 0.09 1/s | 0.09 A/V | 0.065 1/s | (5) |
| $k_{TL}$ | Translation rate | 0.0064 1/s (0.0055 1/s) | 0.0064 A/V ( 0.0055 A/V) | 0.006 1/s | |
| Cp | Ribosome speed | 1.38 nt/s (1.2 nt/s) | 1.38 nt/s (1.2 nt/s) | 2.5 nt/s | |
| $k_{mat}$ | GFP maturation rate | 0.0014 1/s | 0.0014 A/V | 0.000725 1/s | |
| N_RNAP | RNAP consumed per DNA-RNAP during transcription | N = 1 + K_TX * (Lm/Cm) = 2.62 | k = 2.62 | 6.2 | |
| N_Ribo | Ribosome consumed per mRNA-Ribo during translation | N = 1 + K_TL * (Lp/Cp) = 4.33 (N = 4.3) | k = 4.33 (k = 4.3) | 2.92 | |
| [$Ribo_0$] | Total ribosome concentration | 1500 nM ( 850 nM ) | 1100 nV ( 850 nV ) | 1100 nM | |
| $K_{Ribo}$ | Dissociation constant for ribosome binding to RBS on mRNA | 10 nM | R = 1/Km = $10^8$ Ω | 10 nM | |
| *d* | mRNA degradation rate | 0.00091 1/s ( 0.00118 1/s ) | R =1/d =1100 Ω ( R = 850 Ω ) | 0.00083 1/s | |
| $K_i$_TetR | Dissociation constant for TetR dimer and tetO | 5 nM | R= $1/K_i$ = $2*10^8$ Ω | 0.18 nM | (6) |

Note: In our circuits, Michaelis-Menten constants ($K_m$) were used interchangeably with the dissociation constants ($K_d$). All values are the same or comparable to the literature cited with only slight variations for different experimental conditions. **a:** The values in parentheses are parameters used in the modeling of TetR regulation with slight variations to account for different batches/preparations of cell-free lysate and plasmid. **b:** For circuits under a steady-state approximation, capacitors corresponding to binding state variables are removed such that resistors are directly related to steady-state Michaelis-Menten constants ($K_m$) only, and do not affect dynamic parameters like time constants.

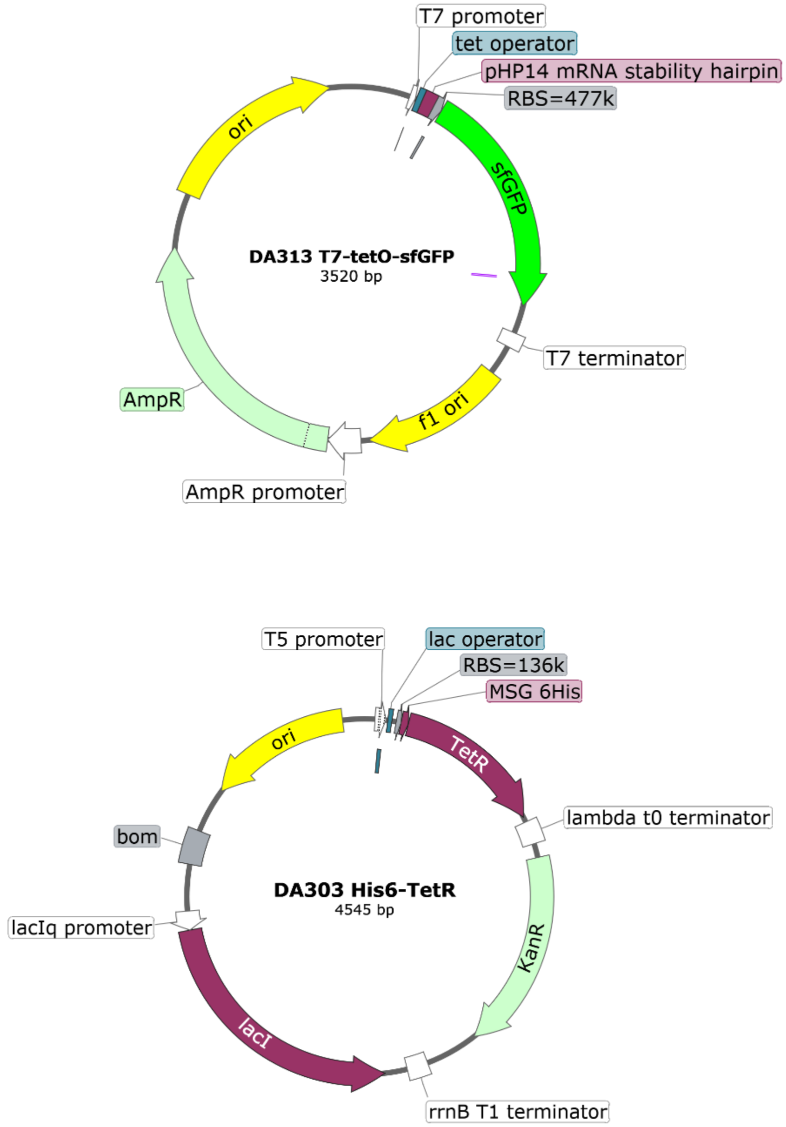


**Figure S2.** Maps of the plasmids used in this work. DA313 is the plasmid used in the cell-free system while DA303 is the plasmid for the preparation of the recombinant TetR protein.

**Supplementary References**


1. Dickinson FM, Dickenson CJ. 1978. Estimation of rate and dissociation constants involving ternary complexes in reactions catalyzed by yeast alcohol dehydrogenase. Biochem J 171:629–37.

2. Ganzhorn AJ, Green DW, Hershey AD, Gould RM, Plapp B V. 1987. Kinetic characterization of yeast alcohol dehydrogenases. Amino acid residue 294 and substrate specificity. J Biol Chem 262:3754–3761.

3. Újvári A, Martin CT. 1996. Thermodynamic and kinetic measurements of promoter binding by T7 RNA polymerase. Biochemistry 35:14574–14582.

4. Skinner GM, Baumann CG, Quinn DM, Molloy JE, Hoggett JG. 2004. Promoter binding, initiation, and elongation by bacteriophage T7 RNA polymerase: A single-molecule view of the transcription cycle. J Biol Chem 279:3239–3244.

5. Marshall R, Noireaux V. 2019. Quantitative modeling of transcription and translation of an all-E. coli cell-free system. Sci Rep 9:11980.

6. Kamionka A, Bogdanska-Urbaniak J, Scholz O, Hillen W. 2004. Two mutations in the tetracycline repressor change the inducer anhydrotetracycline to a corepressor. Nucleic Acids Res 32:842–847.
